# Transient Vascular Cell Integration and Intrinsic Media-Stimulated Endothelial Activation Reveal Diabetic Retinopathy Signatures in Human Retinal Organoids

**DOI:** 10.64898/2026.10.01.755879

**Authors:** Rosalia Tsikandelova, Raphael Schettler, Matthew D Teasdale, Joe Kelk, Veronika Boczonadi, Birthe Dorgau, Rafiqul Hussain, Jonathan Coxhead, Reinhold Medina, Stephen White, Robert Zweigerdt, Lyle Armstrong, Gerrit Hilgen, Majlinda Lako

## Abstract

Retinal organoids (ROs) model human retinogenesis but remain avascular and lack immune components, limiting their utility for studying vascular retinopathies. We generated vascular organoids (VOrs) from human pluripotent stem cells and evaluated their integration into developing ROs. Dissociated VOr cells (day 12 or day 30) were seeded onto 2D retinal neuroepithelium before 3D transition, followed by long-term culture to day 210. VOr-derived cells migrated into ROs and transiently persisted. V30-ROs preserved retinal lamination, major neuronal subtypes and light-evoked retinal ganglion cell responses comparable to non-vascularised controls, whereas media-only ROs cultured in pro-angiogenic conditions showed structural disorganisation and reduced neuronal survival. Single-cell RNA sequencing revealed an endogenous, inflammatory endothelial-like population with partial endothelial-to-mesenchymal features and a perivascular fibroblast-like cluster in pro-angiogenic media-only RO controls. The endothelial-like cell cluster was significantly enriched for genes upregulated in human retinal endothelial cells exposed to high glucose, indicating a diabetic retinopathy–like transcriptional state. IBA1⁺ microglia-like and border-associated macrophage populations were present across all conditions, with developmental shifts in morphology and marker expression. Our findings define conditions under which vascular cells can be incorporated into human ROs without disrupting retinal architecture, while uncovering an intrinsic capacity for inflammatory endothelial activation that partially recapitulates diabetic retinopathy. This work advances the engineering of vascularised, immunocompetent retinal models and highlights endogenous vascular–immune programmes that must be considered when modelling retinal disease *in vitro*.

## Introduction

Vision arises in the retina, the light-sensitive neural tissue at the back of the eye, in which photoreceptors detect photons, interneurons process and refine the signal, and retinal ganglion cells (RGCs) convert it to action potentials that reach higher visual centres of the brain via the optic nerve (Baden *et al*., 2018). Retinal organoids (ROs) derived from pluripotent stem cells are self-organising three-dimensional laminated structures, which reproduce key stages of human retinogenesis marked by the sequential emergence of all retinal neuronal and glial cell types (Dorgau *et al*., 2024). However, conventional retinal organoids (ROs) remain largely avascular and fail to mature beyond a fetal-like developmental stage during prolonged culture (>210 days), as increasing organoid size exacerbates oxygen and nutrient diffusion limitations, ultimately leading to progressive degeneration of RGCs. Existing RO protocols are limited in their ability to model major blinding conditions such as diabetic retinopathy and age-related macular degeneration (AMD), in which inner-retinal barrier failure and autoimmunity are key disease features (Fan and Li, 2025), further highlighting the need for *in vitro* retinal models to incorporate a functional vascular-immune milieu.

To address the need for vascularised and immunocompetent retinal models, several strategies have been employed. The seminal work by Wimmer and colleagues established vascular organoids (VOrs) from human induced pluripotent stem cells (hiPSCs) as a source of pre-organised capillary-like networks with endothelial-pericyte architecture and barrier properties, which can be fused with target organoids to generate vascularised organoids (Wimmer *et al*. 2019). Subsequent protocol adaptations have reproducibly generated hiPSC-derived blood vessels and VOr, providing a robust source of vascular and immune cells that can colonise various organoid types (Meijer *et al*., 2023; Nikolova *et al*., 2025, Sun *et al*., 2022). Taking advantage of these developments, Inagaki *et al*. (2025) were the first to generate primitive, early vascularised ROs (∼day 60) by leveraging VOr and iPSC co-culture without markedly disrupting retinal lamination. The same study reported the integration of endothelial cords and associated pericyte cells, resulting in RO enlargement, increased expression of neuronal markers and improved RGC maturation. Conversely, under diabetic conditions, their vascularised ROs mimicked disease-like responses, which led to a decrease in organoid size and the pervasive loss of RGCs (Inagaki *et al*. 2025).

A recent study by Sharma *et al*. (2025) demonstrated transient RO vascularisation by incorporating ETV2.2-inducible hiPSC-derived endothelial cells (ECs) at the embryoid body (EB) stage. The temporary vascular networks alleviated hypoxia and measurably enhanced RO size, RGC survival and photoreceptor-driven light responses. Despite the favourable effects of endothelial lumens on long-term RGC function, the study reports on the eventual loss of vascular networks past day 150 of retinal development due to the lack of active perfusion (Sharma *et al*., 2025). Although microfluidic systems enhance vascular maturation and survival *in vitro*, achieving long-term vascular stability remains challenging without transplantation of vascularised organoids into a living host (Zhou *et al*. 2025; Kowalski *et al*. 2025). Beyond VOr- or ETV2-based approaches, Abdalkader *et al*. (2026) used a 3D-printed retinal organoid-on-chip device to co-culture Human Umbilical Vein ECs (HUVECs) with ∼day 90 ROs in a fibrin-Matrigel matrix, where ECs organised into RPE-associated vascular networks without perturbing retinal lamination in the short-term (Abdalkader *et al*., 2026).

Despite representing important initial advances toward retinal vascularisation, current vascularised RO models fail to recapitulate the inner blood-retina barrier (iBRB). Instead, they exhibit aberrant vascular patterning, including EC mislocalisation within the normally avascular photoreceptor layer (Sharma et al., 2025), confinement of endothelial cords to the hypoxic core of retinal organoids (Inagaki et al., 2025; Chen et al., 2025), and, in some cases, disruption of overall retinal architecture following vascular integration (Inagaki et al., 2025). Taken together, these limitations underscore the need for a physiologically relevant inner blood–retinal barrier (iBRB), which can contribute to the tissue-specific vascularisation and immune function of the inner retina.

Recent studies have begun to address this gap by incorporating immune components into the developing ROs. Chichagova *et al*. (2023) were among the first to demonstrate the integration of hiPSC-derived microglia-like (iMG) cells into day 90 ROs, showing that immunocompetent organoids can be established without disrupting baseline retinal development. Building on this, Usui-Ouchi *et al*. (2023) provided one of the most anatomically faithful models of RO–microglia integration to date, reporting on the colonisation of plexiform layers in late-stage organoids (∼day 175) by hiPSC-derived macrophage precursors, followed by their gradual maturation into a homeostatic microglial phenotype. Extending these findings, Schmied *et al*. (2024) demonstrated earlier microglial incorporation, with iMG cells integrating into the outer plexiform layer by approximately day 120 via co-culture, within a model of microglia-dependent, ibuprofen-mediated prenatal retinal immune challenge. More recently, Chen *et al*. developed a PDMS microwell-based co-culture system that combines ROs and VOrs to generate more physiologically relevant immunocompetent vascularised retinal organoids Chen *et al*. (2025). This platform improved vascular integration, reduced hypoxia, preserved retinal structure, and supported microglial emergence within the organoid, making it a promising step toward modelling retinal vascularisation and neuroimmune–vascular interactions *in vitro* (Chen *et al*. (2025).

All of the models described above rely on introducing vascular and immune cells into ROs, based on the assumption that these cell types cannot develop alongside neural and retinal lineages because they require markedly different culture conditions. However, IBA1-positive microglia-like cells and border-associated macrophages (BAMs) have been shown to arise intrinsically during early, unguided 2D–3D retinal organoid differentiation, suggesting that mesodermal precursors may be enriched within non-pigmented cystic compartments following low-dose BMP4 supplementation (Bartalska *et al*., 2022).

Despite these advances, the contribution of endogenous mesodermal lineages to vascular and immune compartments in ROs remains poorly defined, and the functional consequences of vascular–immune integration for retinal maturation and disease modelling are unknown. To address these gaps, we generated VOrs from human embryonic stem cells (hESCs) and systematically evaluated their integration into developing ROs using early (day 12) and late (day 30) VOr dissociates, alongside retinal media and retinal-vascular media-only controls, followed by long-term 3D culture until day 210. We show that VOr-derived cells can integrate into ROs and transiently persist, with resulting organoids preserving retinal lamination, major neuronal subtypes and light-evoked RGC responses comparable to non-vascularised controls. Single-cell RNA sequencing further revealed an endogenous, inflammatory endothelial-like population with partial endothelial-to-mesenchymal features and a perivascular fibroblast-like cluster in retinal- vascular media-only RO controls, transcriptionally enriched for high-glucose/diabetic retinopathy signatures. Collectively, our work defines conditions under which vascular and immune cells can be incorporated without disrupting retinal architecture, while uncovering endogenous inflammatory endothelial activation that recapitulates key aspects of diabetic retinopathy *in vitro*.

## Results

### Vascular Organoids Provide a Source of Diverse Vascular and Immune Cells Capable of Retinal Integration

Recent efforts to vascularize ROs using genetically programmable ECs have yielded only transient vascularisation (Sharma *et al*., 2026), failing to recapitulate the physiological emergence of the iBRB and omitting key supporting cell types such as pericytes, smooth muscle cells, astrocytes and glia/microglia. We employed VOrs generated from human pluripotent stem cells to introduce diverse vascular cell types into ROs. Because VOrs naturally contain these essential cell populations, they serve as an ideal platform for vascular integration. VOrs were generated from H9 hESCs using either mechanical agitation (shaking) or stationary culture conditions (**Fig. 1A, B**) following the protocol established by Sun *et al*. (2022). Quantitative real- time PCR analysis demonstrated the downregulation of pluripotency markers (*NANOG, OCT4*) during the course of differentiation and a gradual increase in the expression of astrocytic markers (*GAS6, VIM* and *PRDGFRα*) (**Fig. S1A, B**). Most of the endothelial marker expression reached their peak at D12 (**Fig. S1C**), whilst microglia (*TMEMT11S*, *HEXB*) and pericyte (*PRDGFRβ*) marker expression peaked at D40 (**Fig. S1D, E**). Flow cytometric analysis corroborated these data, demonstrating the highest CD31 cell surface expression at D12 of VOr development (**Fig. S1F- H**).

**Fig. 1:**
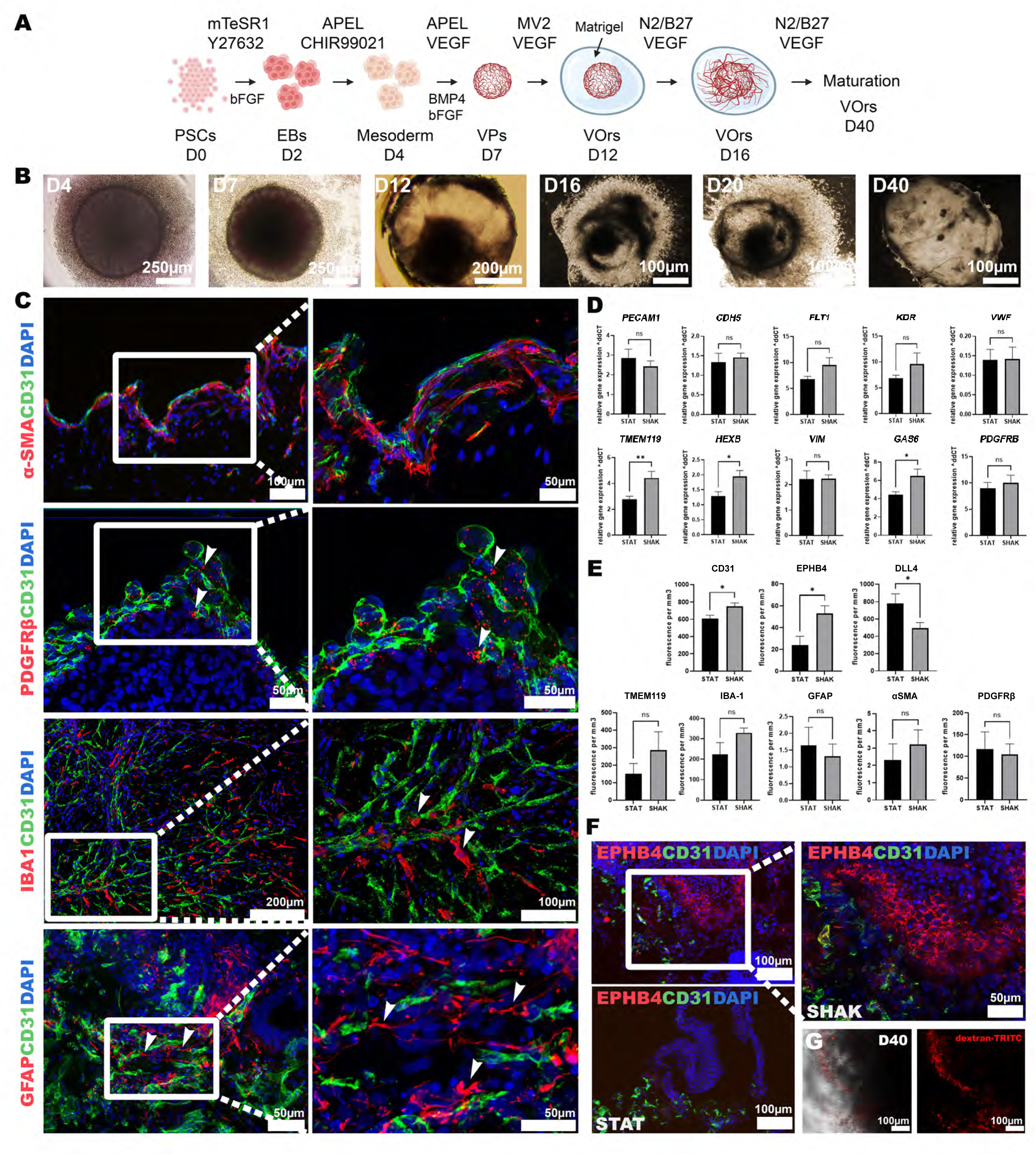
Generation and characterisation of VOrs. **A**. Timeline of VOr differentiation. **B**. Representative bright-field images of VOrs throughout the differentiation procedure, following the darkening of embryoid bodies (D7), cyst formation (D12), sprouting of the cyst following Matrigel embedding (D16 and D20) until its final expansion at D40. **C**. IF Characterisation of D40 VOrs cultured under mechanical agitation for endothelial cell (CD31), smooth muscle cell (α- SMA), pericytes (PDGFR-β), microglia (IBA-1) and astrocytes (GFAP) markers. **D**. Comparison of the relative mRNA levels of *PECAM1, CDH5, VEGFR1, VEGFR2, VWF, TMEM11S, HEXB, VIMENTIN, GASC*, and *PDGFRβ* in D40 VOrs generated under mechanical agitation (SHAK) versus stationary conditions (STAT), normalised to *GAPDH*. Data represent mean ± SEM (n = 3, ***p < 0.001, one-way ANOVA). **E.** IF quantification of vascular marker expression shows a significant increase in CD31 and EPBH4 immunostaining following mechanical agitation. Data represent mean ± SEM (n = 3, ***p < 0.001, two-tailed t-test). **F.** EPBH4 expression is predominantly seen by IF in D40 VOrs cultured under shaking conditions **G.** Dextran-TRITC staining showing tube formation inside VOrs. Nuclei were counterstained with Hoechst.

Immunofluorescence (IF) characterisation of mature day 40 VOrs (D40) cultured under mechanical agitation standard for this protocol described by Sun *et al*. (2022), corroborated the expression of smooth muscle cell marker (α-SMA), pericytes (PDGFR-β), microglia (IBA-1), and astrocytes (GFAP) within close proximity to elongated CD31+ cells (**Fig. 1C**). A qPCR comparison from D40 VORs cultured under shaking versus stationary conditions demonstrated that mechanical agitation led to a significant increase in the expression of microglial (*TMEM119* and *HEXB*) and astrocytic (*GAS6*) markers in D40 VOrs (**Fig. 1D**). IF quantification demonstrated that mechanical agitation increased the abundance of CD31⁺ ECs and enhanced expression of the venous marker EPHB4, while reducing expression of the arterial marker DLL4 (**Fig. 1E, F**). Nevertheless, within shaking-cultured VOrs, CXCR4⁺ arterial ECs were more abundant than CD73⁺ venous ECs (**Fig. S1I**). Further generation of VOrs was carried out by using only mechanical agitation. Notably, day 40 VOrs could uptake dextran-labelled beads, demonstrating the formation of functional, luminal structures capable of internalising exogenous transport molecules (**Fig. 1G**).

To determine if vascular and immune cells within VOrs could integrate into ROs, we generated assembloids by co-culturing two VOrs (D12) with one RO (D90) on Matrigel droplets for 40 days in a 1:1 media mix (Fig. 2A, B) under stationary conditions for four days and subsequently cultured under mechanical agitation. IF analysis revealed that CD31+ cells organised into vessel-like structures and migrated from the VOrs into the basal side of the ROs (Fig. 2Ca). Similarly, GFAP+ astrocytes (Fig. 2Cc-e) and IBA1+ microglia (Fig. 2Cf-k) migrated into the ROs, frequently appearing in proximity to the CD31+ structures. While retinal lamination, indicated by apical photoreceptor precursor (Recoverin) and basal RGC (SNCG) markers, remained intact 14 days post-fusion (Fig. 2Ca, b), this organisation was lost by day 28 onwards (data not shown). These data confirm that VOr-derived vascular and immune cells can integrate into ROs; however, this assembloid approach is ultimately unfeasible for maintaining a laminated, vascularised, and immunocompetent retinal model in the long term.

**Fig. 2:**
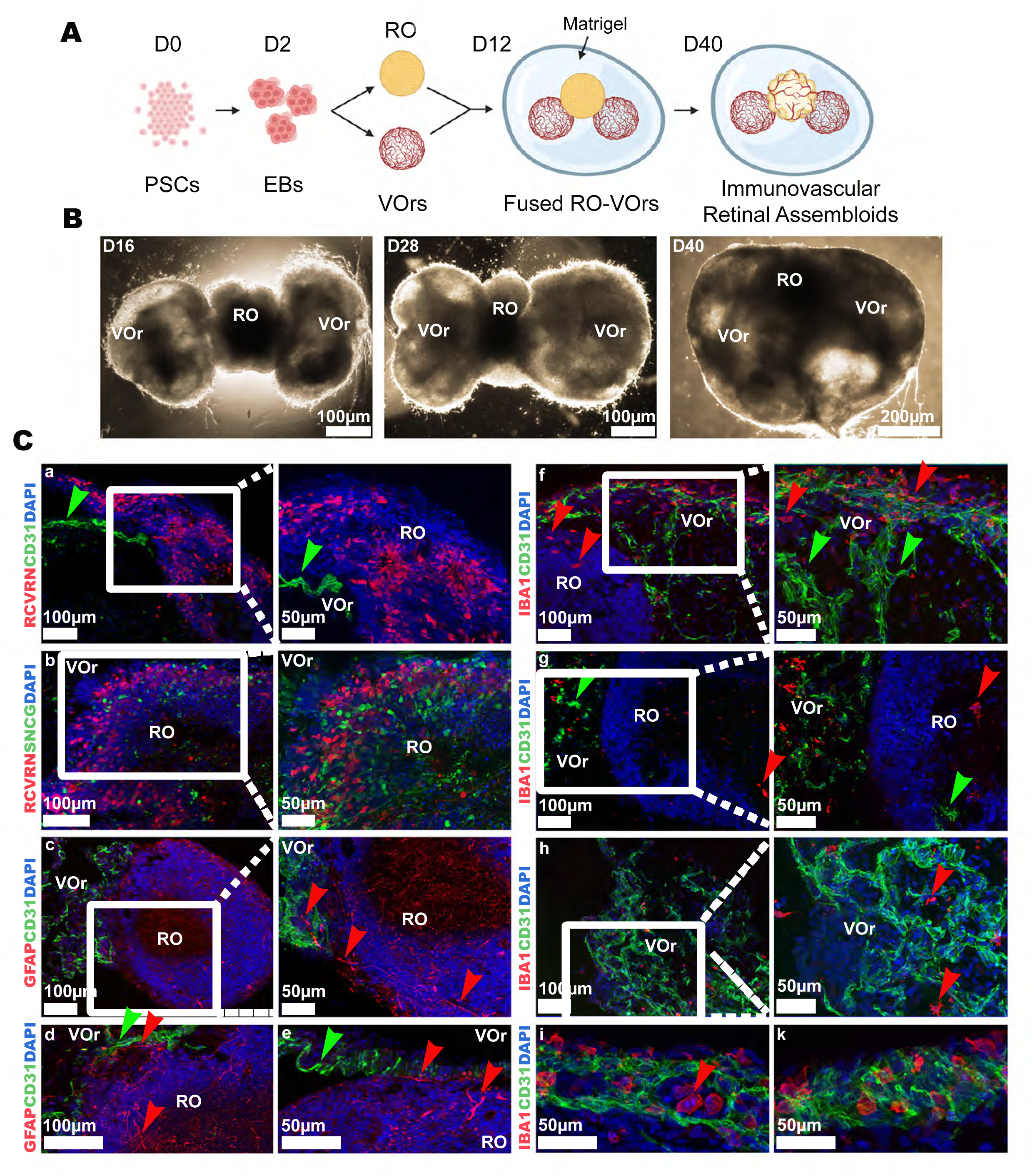
Assembloid formation of DG0 ROs with D12 VOrs enables the incorporation of ECs, astrocytes and microglia into the retinal compartment. **A.** Schematic briefly outlines the generation of assembloids from D90 R0s and D12 VOrs. **B.** Brightfield images show assembloid formation at 4 days (D16), 16 days (D28) and 28 days (D40) after Matrigel embedding-facilitated fusion. C. *a*) Top left panels show an example of CD31+vessel-like (green arrows) within the RO, with the same RO showing expression of pan photoreceptor maker Recoverin (RCVRN). *b*) The panel below depicts the retinal compartment of a fusion with relatively good preservation of retinal lamination, marked by RCVRN+ photoreceptors apically and SNCG+ ganglion cells on the basal aspect. *c-d*) Bottom left panel images show GFAP+ astrocytes migrating from VOr into ROs (red arrows), whereas CD31+ vessel-like structures are indicated by green arrows. *f*) The top right panel shows IBA+ microglia (red arrows) and CD31+ cells (green arrows) within a VOr compartment of a fusion. In contrast, the image below (*g*) depicts the invasion of IBA+ (red arrows) and CD31+ cells from the VOr compartment into the ROs. *h*) VOr section showing CD31+ endothelial and IBA-1+ microglial-like cells. *i*) The bottom-right panel image displays a selected vessel-like region from assembloids with IBA+ cells residing and lining the inside of a CD31+ vessel-like structure. Nuclei were counterstained with Hoechst.

## VOr-Derived Cell Integration and Survival in 3D Retinal Organoids

Recent studies have shown that co-culturing ECs with developing 2D retinal neuroepithelium, followed by 3D transition, results in transient RO vascularisation (Sharma *et al*., 2026), suggesting that early-stage integration is feasible. To identify optimal conditions for the co-survival and integration of endothelial-, pericyte-, and microglia-derived VOrs within ROs, we evaluated several media formulations. These included cell type-specific media, standard retinal differentiation media (RDMI) and a 1:1 mixture of RDMI and endothelial growth media (EGM2), both with and without VEGF supplementation (**Fig. S2A**).

Using Endothelial Colony-Forming Cells (ECFCs) as a proxy for primary vasculature, we found that the 1:1 RDMI: EGM2 mix supplemented with VEGF most closely recapitulated the performance of standard endothelial EGM2 medium and enhanced cell survival relative to all other tested conditions (**Fig. S2A**). In contrast, HMC3 microglia and human brain microvascular pericytes exhibited lower sensitivity to media composition, maintaining high viability across all tested groups.

Based on these findings, we designed a four-arm experimental framework (**Fig. S2B**). The Retinal Control (**RC**) group followed the established 2D-3D protocol (West *et al*. 2022) without vascular/immune cell inclusion or media modification. The **RC-V** group served as a media control, utilising the RDMI: EGM2+VEGF formulation without VOr cellular integration. To leverage the temporal emergence of cell types within VOrs, where pericytes appear early, followed by ECs, smooth muscle, and microglia, we established two experimental groups (Sun *et al*. 2022). These involved the dissociation of either early-stage (D12, **V12-RO**) or late-stage (D30, **V30-RO**) VOrs, which were then plated onto 2D developing retinal neuroepithelium generated from CRX-GFP hESCs at D30 (Collin *et al*. 2016), followed by lifting of retinal structures (D44) and culture in 3D until day 210 of differentiation (**Fig. S2B**). To track the spatial localisation of vascular cells, a parallel experiment was conducted in which mCherry+ VOrs were dissociated and added to the developing neuroepithelium, followed by bright-field imaging at key stages of differentiation (**Fig. S3A**). mCherry+ cells were observed in proximity to the developing optic vesicles under 2D culture conditions. Following the excision of retinal structures and subsequent 3D culture, mCherry+ cells localised to the putative ciliary margin, developing retinal neuroepithelium and the basal side of ROs. Quantification revealed that the proportion of ROs containing mCherry+ vascular cells remained high in both V12- and V30-RO groups until day 60, when a sharp decline occurred. This was followed by a gradual loss of vascular cells over time (**Fig. S3B**), ultimately leaving 10% of V12-ROs and 30% of V30-ROs with detectable exogenous vascular cells by day 90 of differentiation. By day 100 of differentiation, no mCherry+ vascular cells could be detected within the V12- or V30-ROs.

By day 210, all experimental groups produced mature organoids featuring distinct outer segments (**Fig. S2C, D**). However, differentiation efficiency varied significantly across conditions: the RC group was the most efficient (93.5%), followed by V30-RO (83.6%) and V12- RO (50%), while the RC-V media control exhibited the lowest efficiency (25.2%). These results suggest that while the 1:1 retinal: endothelial media mix supplemented with VEGF negatively impacts standard RO development, the incorporation of VOr-derived cells, even transiently, appears to partially mitigate these adverse effects.

## Effects of Transient Vascular Cell Integration on Retinal Organoid Development

To evaluate how the integration of vascular cells, albeit transiently, influences RO differentiation and photoreceptor maturation, we utilised IF and high-density microelectrode array (MEA) recordings. By day 60, the RC group exhibited highly organised ROs characterised by distinct apical-basal polarity: VSX2+ retinal progenitor cells were localised to the apical side, while SNCG+ RGCs were localised basally (**Fig. 3a**). Notably, radial-like CD31 expression was observed at days 60 and 90 of differentiation (**Fig. 3b-e**). Further investigation revealed that this radial CD31 expression did not co-localise with Müller glial (vimentin) or astrocytic (GFAP) markers. Instead, CD31 partially co-localised with ATOH7, a marker of neurogenic progenitors identified through single-cell RNA-sequencing analysis (Sridhar *et al*. 2020; Dorgau *et al*. 2024). By day 90, Recoverin+ photoreceptor precursors emerged apically (**Fig. 3f**); however, the absence of endothelial lectin UEA-1 staining at this time point indicated that no ECs were present.

**Fig. 3.**
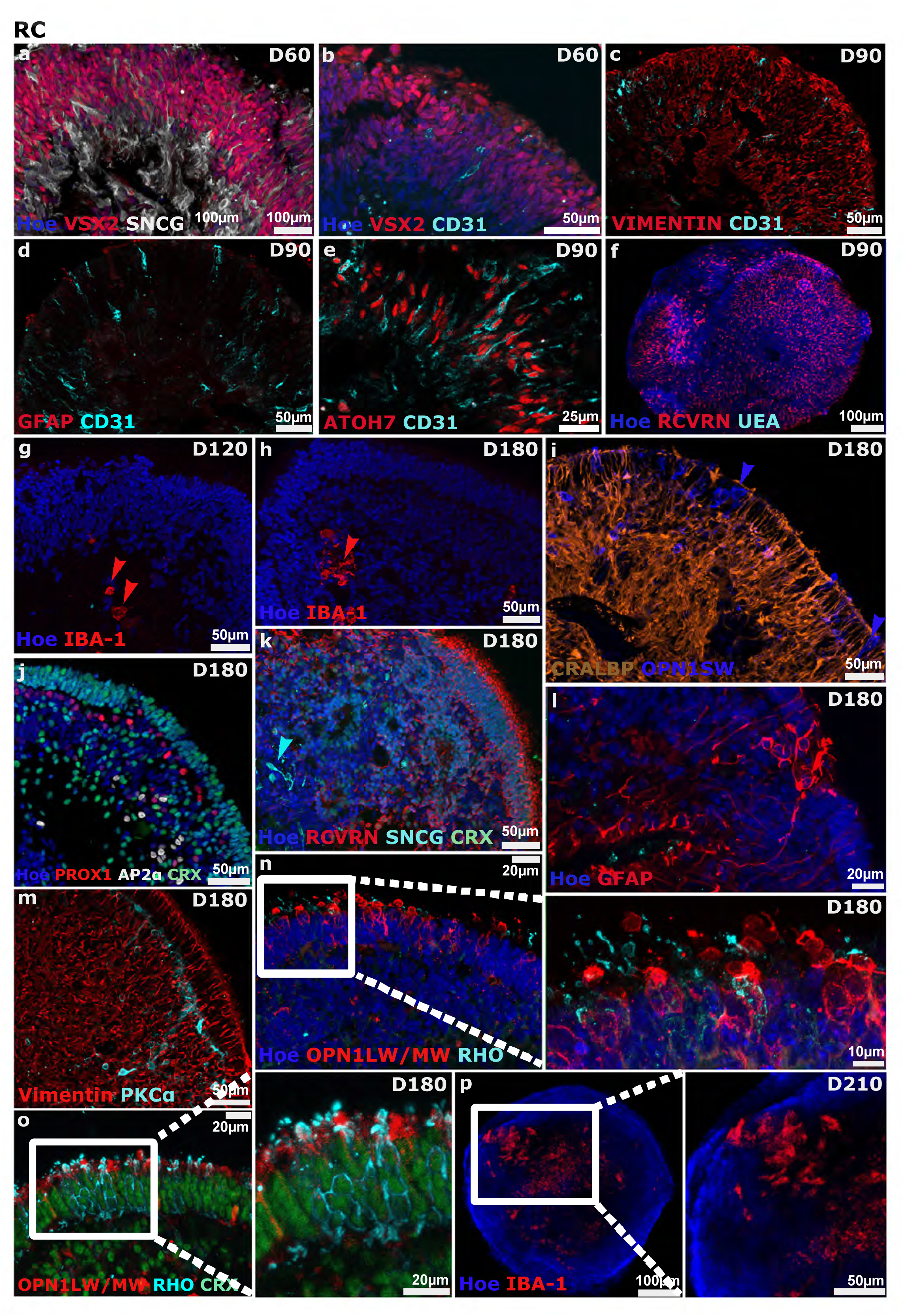
Immunofluorescence characterisation identifies putative microglia-like IBA1⁺ cells in control retinal organoids at D120, 180 and 210 of differentiation. a-b. At D60, control CRX- GFP+ROs stained for VSX2⁺ retinal progenitor cells (red) and SNCG⁺ retinal ganglion cells (white). CD31⁺ cells (light blue) were radially positioned within the apical region of the ROs. **c-e.** In D90 ROs, CD31 co-localised with ATOH7⁺ neurogenic progenitor marker but not with Müller glia or astrocytic markers (vimentin and GFAP, respectively). **f.** D90 whole-mounts showed no staining for vascular endothelium by UEA-I (light blue) on confocal imaging. **g-h.** IBA-1+ staining (red) was observed in D120 and D180 control ROs. At D180, ROs showed expression for Müller glia (CRALBP, orange), short wavelength cone photoreceptors (OPN1SW, blue) **i**, horizontal (PROX1, red) and amacrine cells (AP2α, white) **j,** photoreceptors (RCVRN, red) and ganglion cells (SNCG, light blue) **k,** astrocytes (GFAP, red), Müller glia (vimentin, red) and bipolar cells (PKCα, light blue) **l-m**, as well as mature photoreceptor markers (OPN1LW/MW, red; Rhodopsin, light blue) **n, o**. **p.** Whole-mount clearance shows the retention of IBA-1 expression at day 210. CRX-GFP marks retinal photoreceptors (green). Nuclei were counterstained with Hoechst.

In contrast, IBA-1+ immune cells were detected within the ROs as early as day 120 and persisted until day 210 (**Fig. 3g, h, p**). By day 180, RC ROs reached developmental maturity, displaying well-defined retinal lamination and the presence of all major retinal cell types. This maturity was confirmed by the detection of PROX1+ and AP2α+ horizontal and amacrine cells, as well as sparse SNCG+ RGCs (**Fig. 3j, k**). The presence of Müller glia was evidenced by CRALBP and Vimentin staining (**Fig. 3i, m**), while PKCα+ bipolar cells were situated adjacent to apically aligned mature cones and rods, marked by OPN1LW/MW, OPN1SW, and Rhodopsin immunostaining (**Fig. 3i, m-o**). Additionally, we identified GFAP+ cells with morphologies distinct from Müller glia (**Fig. 3l**). These likely represent endogenous astrocytes, consistent with previous findings from our laboratory (Dorgau *et al*. 2022). Confocal immunostaining of cleared ROs from the RC group demonstrated the presence of Iba-1+ cells (**Fig. 3p**) inside the organoids.

Similarly, RC-V ROs exhibited distinct lamination up to day 120 of differentiation (**Fig. 4a-f**) and a radial-like CD31 expression, which partially co-localised with neurogenic progenitor marker ATOH7, but not GFAP or Vimentin (**Fig. 4c-e**). Developing photoreceptors marked by Recoverin and CRX expression were localised at the apical aspect of ROs up to day 120 of differentiation (**Fig. 4f-i**). However, this organisation was lost by day 180, characterised by the basal migration of photoreceptors (**Fig. 4i**) and the apical displacement of RGCs (**Fig. 4l**). There was also a marked disorganised location of horizontal and amacrine cells (PROX1+, AP2α+) and atypical Müller glia immunostaining on the apical surface (**Fig. 4m, n)**. This structural disorganisation was further corroborated by the presence of rod photoreceptors on the basal aspect of RC-V ROs (**Fig. 4p**). In contrast to RC ROs, the RC-V ROs displayed the presence of microglia-like cells mostly during late-stage differentiation (day 180; **Fig. 4g, k**), alongside increased GFAP+ astrocytic immunostaining (**Fig. 4o, q**), exemplified by GFAP and PDGFRα immunostaining. Despite not adding any VOr cells to this group, a few CD31+ endothelial-like cells were detected as early as day 90 (**Fig. 4f**) by UEA staining and persisted until the late differentiation stages, often forming clusters within the organoid core (**Fig. 4i, s**). Furthermore, cells expressing the smooth-muscle marker αSMA were also observed (**Fig. 4t**). Collectively, these data suggest an intrinsic capacity for mesodermal differentiation within ROs, likely driven by the combination of retinal and endothelial media supplemented with VEGF.

**Fig. 4.**
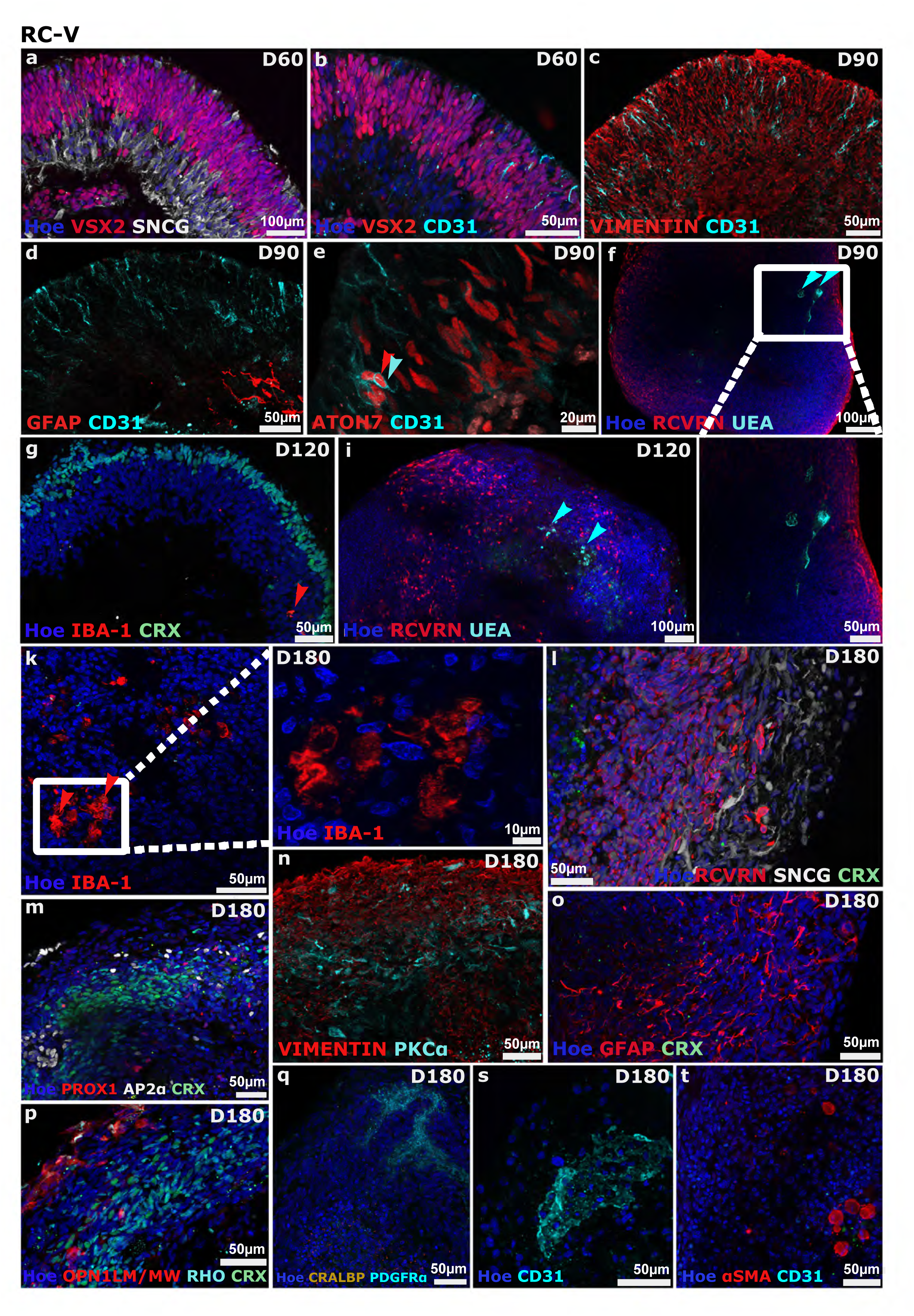
Immunofluorescence reveals endogenous vascular clusters in RC-V retinal organoids and a substantial loss of retinal lamination by D180. **a.** At D60, RC-V ROs exhibit laminar architecture as shown by the apical expression of VSX2+ retinal progenitors (red) and the basal positioning of underlying SNCG+ ganglion cells (white). **b.** Staining shows radial expression of CD31+ (light blue) adjacent to VSX2+ retinal progenitors (red). **c-e.** Transient CD31+ expression (light blue arrows) in RC-V ROs partially co-localises with neurogenic ATOH7+ progenitors, but not Müller glia (Vimentin, red) or astrocytes (GFAP, red) at D90. **f.** Confocal imaging reveals sparse endothelial-like structures labelled with UEA-I (light blue arrows) **g.** Emergence of IBA1⁺ cells in RC-V medium-control ROs by day 120 (red arrow)**. i.** Staining confirms the presence of small clusters of UEA⁺ endothelial cells in day-120 RC-V ROs (red arrow)**. k.** Day-180 RC-V ROs contained IBA1⁺ cells (red) displaying both amoeboid and ramified morphologies. **i.** IF at D180 shows severely disrupted retinal lamination, as evidenced by the inverted expression of SNCG+ ganglion cells (white) and RCVRN+ photoreceptors (red). **m-p.** Characterisation of mature D180 RC-V ROs shows aberrant expression patterns for most retinal markers, including PROX1 (horizontal cells) and AP2α (amacrine cells), vimentin (Müller glia), PKCα (bipolar cells), GFAP (astrocytes), OPN1LM/MW (long- and medium-wavelength cones) and Rhodopsin (rods). **q.** D180 RC-V ROs showed loss of Müller glia (CRALBP, orange) and the presence of PDGFRα+ astrocytes (light blue)**. s, t.** IF shows clusters of endogenous CD31+ ECs (light blue) and αSMA+ smooth muscle cells (red). CRX-GFP marks retinal photoreceptors (green). Nuclei were counterstained with Hoechst.

Despite being cultured in the same media as the RC-V group, V12- and V30-ROs maintained robust retinal lamination throughout differentiation. Although V12-ROs exhibited transient rosette-like structures at day 60 (**Fig. 5a**), prominent CD31+ cells organised either centrally or in a radial fashion were noted at days 60 and 90 (**Fig. 5b, c-e**). The radial-like pattern of CD31 expression was similar to that observed in the RC and RC-V groups, with partial co-localisation with the neurogenic progenitor marker ATOH7, but not with GFAP or Vimentin (**Fig. 5c–e**, **Fig. 6b- e**). In contrast, the central localisation of CD31 expression in V12-ROs and V30-ROs is more consistent with the presence of endothelial-like cells within the organoids. This interpretation was supported by staining with the endothelial lectin UEA at day 90 (**Fig. 5f**; **Fig. 6f**). While CD31+ cells persisted until the end of the culture period, their abundance decreased after day 90 (**Fig. 5h**; **Fig. 6l**), corroborating the transient integration of VOr data revealed by mCherry+ cells in **Fig. S3**. Notably, in V12-ROs, CD31+ endothelial-like cells organised into vessel-like structures adjacent to the photoreceptors (**Fig. 5m**), GFAP+ astrocytes (**Fig. 5p)**, or smooth muscle-like cells (**Fig. 5m**). Cord-like CD31+ cells were also found in close proximity to Iba1+ cells by confocal microscopy in both V12- and V30-ROs (**Fig. 5q**, **Fig. 6p**). The presence of vascular-like lumens lined up with putative ECs and pericytes could also be seen at day 210 in RC-V, V12- and V30-ROs on transmission electron microscopy, whereas RC-V ROs displayed a high abundance of RPE cells (**Fig. S4**).

**Fig. 5.**
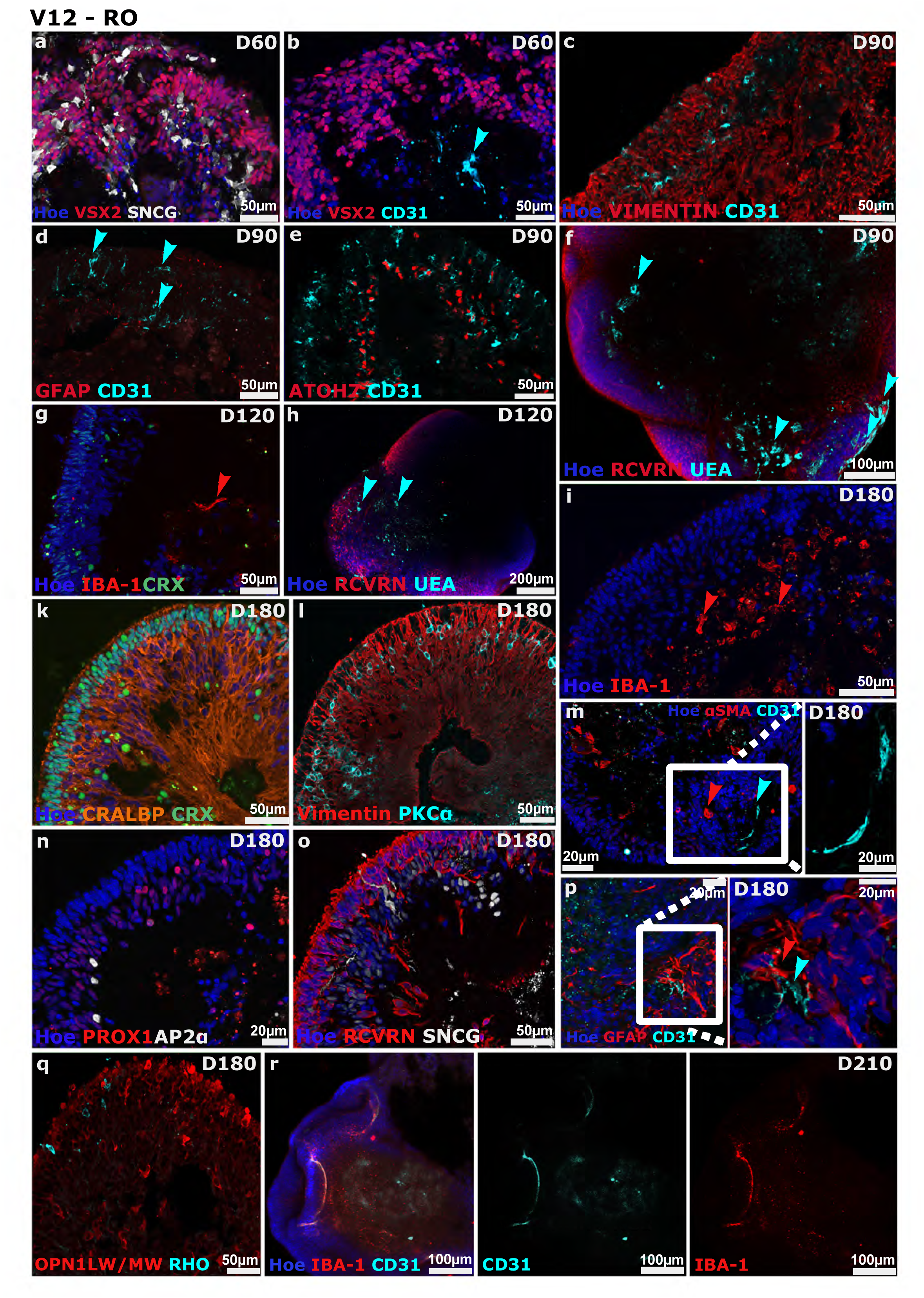
V12-RO development is consistent with the long-term preservation of retinal structure and the presence of vascular cells between DG0 and D210 of differentiation. **a.** At D60, V12-ROs show disorganised expression of VSX2+ retinal progenitors and SNCG+ ganglion cells. **b.** CD31+ cells can be seen at the basal side of V12-ROs at day D60 (light blue arrow). **c-e.** CD31expression is observed radially in D90 V12-ROs (light blue arrows), partially co-localising with ATOH7+ neurogenic progenitors but not Vimentin+ Müller glia or GFAP+ astrocytes**. f.** Whole-mount confocal imaging of V12-ROs at D90 shows RCVN+ photoreceptors and corroborates endothelial cell presence via UEA-I staining (light blue arrows). **g, h**. V12-ROs present with ramified IBA-1+ microglial-like cells (red arrow) and UEA-I+ endothelial cells (light blue arrows) at D120. **i.** IF shows prevalent expression of IBA-1+ microglial-like cells in V12-ROs at D180 (red arrows). **k-l.** Immunostaining shows retinal-characteristic expression of CRALBP+ (orange) and vimentin+ (red) Müller glia, as well as PKCα+ bipolar cells (light blue) in the inner nuclear layer of V12-ROs at D180. **m-p.** D180 V12-ROs are present alongside CD31+ ECs (light blue arrow) near αSMA+ smooth muscle cells (red arrow) and GFAP+ astrocytes. D180 V12-ROs stained for PROX1+ horizontal and AP2α+ amacrine cells, as well as RCVRN+ photoreceptors and SNCG+ ganglion cells. **q.** IF corroborates V12-RO maturation by the expression of long- and medium-wavelength cones (OPN1LW/MW - red) and rod (Rhodopsin- light blue) photoreceptor markers, respectively, at D180. **r.** Confocal imaging of D210 V12-RO wholemounts revealed cord-like expression of closely associated IBA-1+ and CD31+ cells. CRX-GFP marks retinal photoreceptors (green). Nuclei were counterstained with Hoechst.

**Fig. 6.**
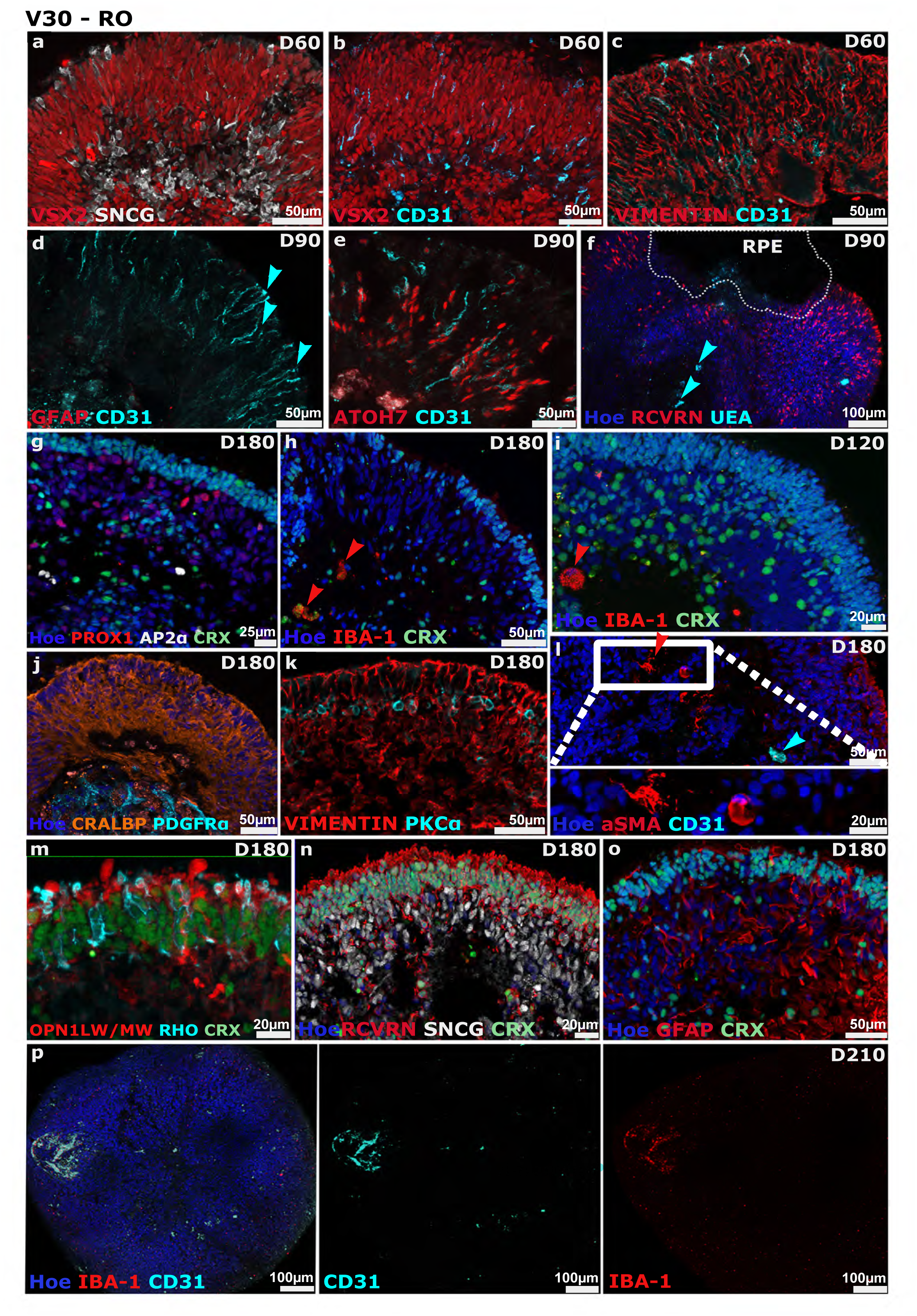
V30-ROs maintain retinal architecture by immunofluorescence throughout the course of retinal differentiation. **a.** IF depicts normal early organoid development marked by the expression of VSX2+ retinal progenitors (red) and underlying SNCG+ ganglion cells (white). **b.** Transient CD31+ radial expression (light blue) at D60 is nestled between VSX2+ progenitors. **c-e.** Radial CD31 expression (light blue) partially co-localises with ATOH7, a marker of neurogenic progenitors (red), but not Vimentin+ Müller glia or GFAP+ astrocytes (red). **f.** Confocal image shows staining of RCVRN+ photoreceptors and UEA-I+ vascular endothelium at D90. **g.** At D180, V30-ROs exhibit layering of PROX1+ horizontal (red) and AP2α+ cells (white). **h.** IF shows positive staining for IBA-1+ cells (red arrows). **i.** Confocal image shows staining of RCVRN+ photoreceptors and UEA-I+ vascular endothelium. **j, k.** D180 V30-ROs staining for Müller glia (CRALBP, orange; Vimentin, red) and astrocytes (PDGFRα, light blue), and bipolar cells (PKCα, light blue). **l.** D180 staining confirms the presence of αSMA+ smooth muscle cells (red) as well as CD31+ endothelial cells (light blue) in V30-ROs. **m-o.** V30-ROs exhibit retinal maturation, as evidenced by the expression of RCVRN+ photoreceptors (red), SNCG+ ganglion cells (white), long- and medium-wavelength cones (OPN1LW/MW, red), rods (light blue), and GFAP+ astrocytes by D180. **p.** Confocal imaging of cleared whole mounts reveals close association of CD31+ and IBA-1+ cells within a putative vascular structure at D210. CRX–GFP marked retinal photoreceptors (green). Nuclei were counterstained with Hoechst.

Unlike the RC and RC-V ROs, IBA-1+ microglia-like cells were integrated much earlier, by day 60, and persisted into late stages (day 180–210) (**Fig. 5g, i**; **Fig. 6h, i)**, where they frequently co-localised with CD31+ structures (**Fig. 5r**; **Fig. 6p**). Furthermore, both V12- and V30-ROs contained GFAP+/PDGFRα+ astrocyte-like cells with distribution patterns distinct from CRALBP+ Müller glia (**Fig. 5p**; **Fig. 6j**). Crucially, all major retinal markers were detected at their correct anatomical positions: photoreceptors were aligned at the apical edge (**Fig. 5k, o, q**; **Fig. 6m-o**) in proximity to bipolar cells (**Fig. 5l**; **Fig. 6k**), horizontal and amacrine cells were distributed within the putative inner nuclear layer (**Fig. 5n**; **Fig. 6g**), and RGCs were correctly localised to the basal aspect (**Fig. 5o**; **Fig 6n**).

To evaluate the functional impact of transient vascular cell integration on retinal circuitry, we recorded RGC activity using high-density multielectrode arrays (MEAs) with 4096 recording electrodes. Following established protocols (Hallam *et al*. 2018), organoids were flattened onto the MEA with the RGC layer in direct contact with the electrodes. Because RGC activity integrates the vertical signalling pathway from photoreceptor phototransduction over bipolar cell transmission to RGC spiking activity, MEA recordings serve as a comprehensive readout of global inner and outer retinal health, as well as synaptic integrity. All ROs were stimulated with a range of light stimuli (full-field flashes and gratings) at varying contrasts to assess functional maturity and integrity. Due to the loss of structural lamination in the RC-V group, functional comparisons were restricted to the RC, V12-, and V30-RO cohorts. While V12-ROs exhibited a significant decline in light-evoked increased activity compared to the RC group, the V30-ROs performed comparably to the RC controls (**Fig. S5A, D**), suggesting that vascular cell integration in the V30 condition does not impede physiological light responses.

To confirm that recorded light responses were driven by photoreceptor phototransduction rather than intrinsically photosensitive RGCs, we utilised 8-br-cGMP puffs to mimic the "dark current." In the dark, cGMP keeps photoreceptor CNG channels open, which allows sodium and calcium to enter and depolarise the photoreceptors. The binding of cGMP should be depicted as a change in spiking, and RGCs exhibiting decreased spiking are categorised as photoreceptor-driven ON-type cells. We observed no significant differences across the three groups regarding reduced firing rates in response to cGMP (**Fig. S5E**). Similarly, no significant disparities were found in cells showing increased firing rates, presumably representing OFF-type RGCs (**Fig. S5B**). We further assessed the maturation of functional neural networks by measuring responses to gamma-aminobutyric acid (GABA), the main inhibitory transmitter in the CNS. All three RO types exhibited comparable responses to GABA puffs, with no significant differences in either increased or decreased firing rates (**Fig. S5C**, **F**). Taken together, these functional data demonstrate that the transient integration of vascular cells in the V30-ROs preserves the integrity of the vertical signalling pathway and maintains the maturation of neural networks, resulting in light-responsive organoids that are physiologically comparable to non-vascularized controls.

## Phenotypic Characterisation of Endogenous Microglia-Like and Border-Associated Macrophage Populations in Retinal Organoids

Given the pervasive emergence of IBA-1+ cells throughout all experimental groups until day 210 (**Fig. 3-6**), we probed IBA-1 expression by IF in combination with other microglial-related markers to determine whether IBA-1+ cells represent a *bona fide* microglial population in our ROs (**Fig. S6**). All groups displayed both ameboid and more ramified IBA1⁺ morphologies by IF, which could pertain to a variety of microglial states as well as closely associated myeloid cell types. Partial co-localisation of IBA-1 immunostaining with BAM marker CD163 at day 180 in all experimental groups suggested that a subset of the mature IBA-1+ cells, may represent BAMs, which could nevertheless be phagocytic in nature (**Fig. S6A-D**). Consistent with a more resident microglial phenotype, we observed co-localisation of the homeostatic microglial marker P2RY12 with the microglia-specific marker TREM2 in V12- and V30-ROs at day 120 (**Fig. S6C, D**). P2RY12⁺/TMEM119⁺ putative microglia-like cells were also detected by IF in RC-ROs at day 180 (**Fig. S6A**). Expression of P2RY12 and TMEM119 microglia markers at both timepoints was accompanied by predominantly ameboid morphologies, indicating that these resident microglia-like cells can adopt activated-like states despite their homeostatic marker profile. By day 180, V12- and V30-ROs had lost detectable P2RY12 expression, in contrast to RC-ROs, which retained P2RY12 but no longer showed co-localisation with TMEM119 (**Fig. S6A, CD, D**). The peak of reactive morphologies at day 120 corresponded with an active remodelling phase of synaptic pruning, which may account for the higher incidence of ameboid microglia, prior to the acquisition of more “settled” ramified phenotypes in RC-ROs at day 180. Notably, this timing is consistent with recent work showing that exogenously added microglia incorporate most efficiently into ROs once retinal lamination is largely established by day 120 (Schmied *et al*., 2025). Although P2RY12⁺/TMEM119⁺ co-expression is considered highly specific for microglia, neither IBA1, P2RY12 nor TMEM119 is exclusively expressed by all microglial subtypes (Kenkhuis *et al*., 2022). Hence, the pronounced increase in P2RY12⁺/TMEM119⁺ cells at day 120 likely reflects a transient, transitioning microglia-like state rather than a stable, homogeneous population.

## Endogenous inflammatory endothelial activation in RC-V ROs recapitulates diabetic retinopathy

To analyse the cellular composition of the day 210 ROs, we performed single-cell RNA sequencing (scRNA-seq) across all four experimental groups. Following rigorous quality control, the resulting transcriptomes from 22,933 cells were sequenced, with an average of 42,464 reads and 1,843 genes per cell, were integrated and clustered using Seurat based on highly variable genes (**Table S1;** **Fig. 7A**). This analysis revealed 26 distinct clusters encompassing all major retinal lineages, including rod and cone photoreceptors, horizontal and amacrine cells, bipolar cells, RGCs, Müller glia, and RPE cells. While RGCs were observed at low frequency in the control RC-ROs, their presence was further reduced in all three groups of ROs (RC-V, V12-RO and V30- RO) cultured with retinal-endothelial media supplemented with VEGF (**Fig. 7B**). Notably, we identified a distinct astrocyte cluster characterised by high *GFAP* expression, which corroborated our immunofluorescence findings. Although this population was present in RC- ROs and RC-V ROs, it was enriched in V12- and V30-ROs (Fig. 7B). This enrichment may reflect the contribution of astrocyte-like cells from the VOrs and/or vascular-mediated expansion of the endogenous astrocyte population. However, lineage-tracing experiments will be required to distinguish between these possibilities.

**Fig 7:**
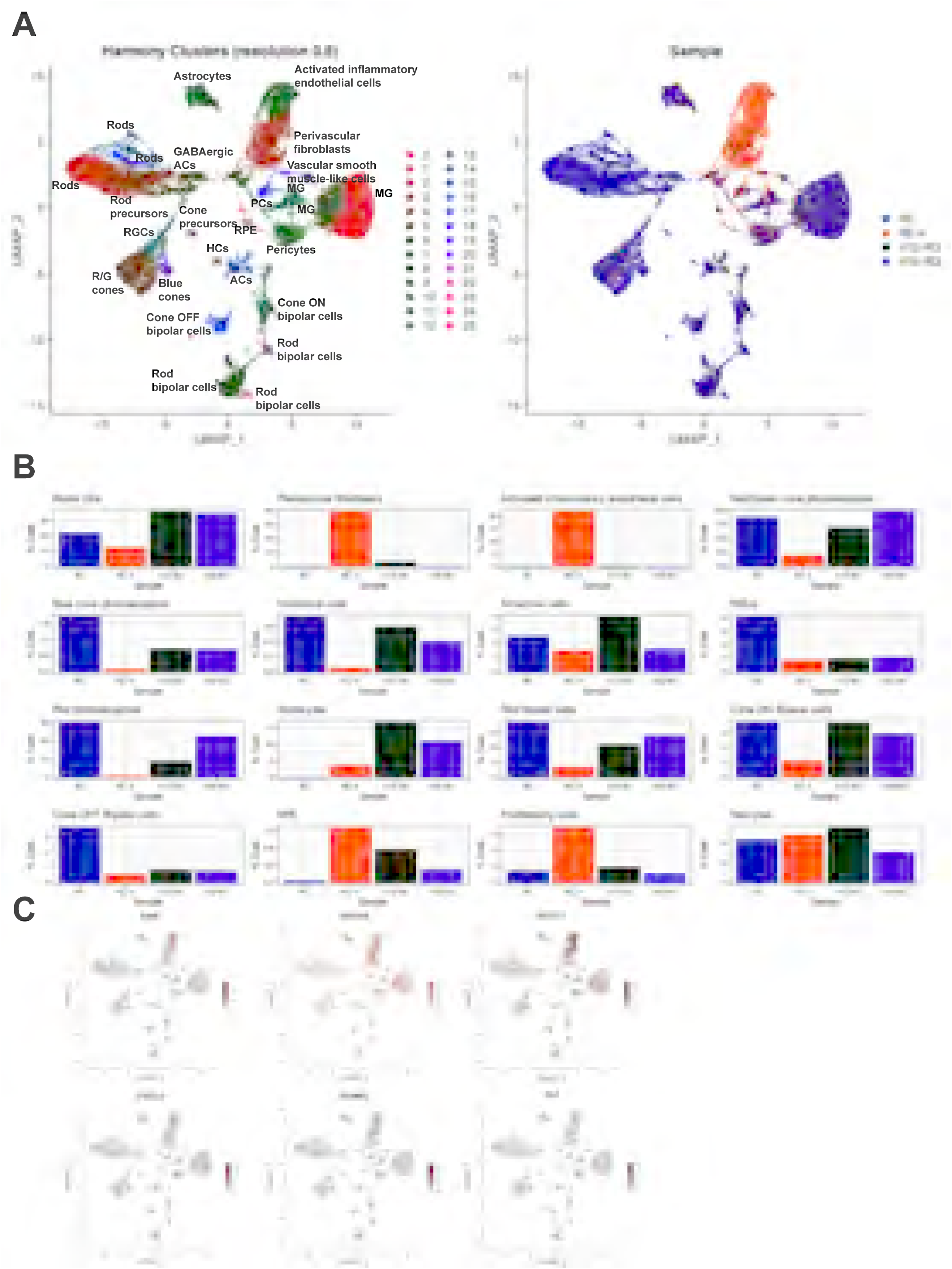
Single-cell RNA sequencing (scRNA-seq) characterisation and cell proportion analysis of retinal and supportive cell populations. **A.** UMAP dimensional reduction plots showing single-cell clustering. *Top left:* Distribution of cells categorised by sample condition (RC, RC-V, V12-RO, V30-RO). *Bottom left:* Unsupervised cell clustering identified at a resolution of 0.8 (clusters 0–25) and annotated by key retinal and vascular/stromal cell types, including Rods, Red/Green cones, Blue cones, Rod/Cone precursors, Retinal Ganglion Cells (RGCs), Amacrine Cells (ACs), Horizontal Cells (HCs), Bipolar Cells (Cone ON, Cone OFF, Rod), Müller Glia (MG), Astrocytes, Pericytes, Activated Inflammatory ECs, Perivascular Fibroblasts, Vascular Smooth Muscle-like Cells, and Retinal Pigment Epithelium (RPE). **B.** Quantification of cell population proportions (% Cells) across experimental conditions (RC, RC-V, V12-RO, and V30-RO) across identified cell types, including Müller Glia, Blue Cone Photoreceptors, Rod Photoreceptors, Cone OFF Bipolar Cells, Perivascular Fibroblasts, Horizontal Cells, Astrocytes, RPE, Activated Inflammatory ECs, Amacrine Cells, Rod Bipolar Cells, Proliferating Cells, Red/Green Cone Photoreceptors, RGCs, Cone ON Bipolar Cells, and Pericytes. **C.** Feature UMAP plots displaying spatial expression levels of marker genes associated with vascular, endothelial, and inflammatory signalling pathways, including *EGFL7*, *VEGFA*, *ENG*, *IL6*, *ICAM1*, and *CXCL2*. Relative expression intensity is indicated by a colour scale.

Consistent with Sharma *et al*., our scRNA-seq data did not identify a cluster with a canonical endothelial transcriptional signature (e.g., *CDH5, KLF2, VWF, PECAM1*), despite the detection of CD31⁺ cells by IF. However, we identified a prominent population (cluster 8), particularly enriched in RC-V ROs (**Fig. 7A, B**). Although canonical endothelial markers, including *PECAM1* and *KDR,* were not detected, residual endothelial identity was supported by the expression of *CDH5, EMCN, ENG, EGFL7, ECSCR,* and *CLEC14A* (**Fig. 7C, Table S1**). This cluster expressed markers associated with activated, inflammatory endothelial-like cells, including chemokines (*CXCL2*), cytokines (*IL6*), angiogenesis-related genes (*VEGFA, EGFL7*), and inflammatory mediators (*ICAM1*). Notably, these cells retained moderate expression of *ENG* (endoglin), a marker present in both quiescent and activated endothelial states (**Fig. 7C, Table S1**). This cluster also exhibited features consistent with partial endothelial-to-mesenchymal transition (EndMT) (Piera-Velazquez and Jimenez, 2019) exemplified by the concurrent expression of mesenchymal and extracellular-matrix-associated genes, including *ACTA2, FN1, COL1A1, COL5A2, COLCA2, COLCA3, POSTN, VCAN, CTHRC1, SERPINE1, MMP11, MMP14*, and *SNAI1* (**Fig. S7**). However, the absence of a comprehensive contractile mesenchymal programme, including *TAGLN, CNN1, MYH11, VIM* and *CDH2*, together with the persistence of endothelial transcripts, argues against a complete EndMT (**Fig. S7**). Therefore, this population was interpreted as comprising partially transitioned ECs with an EndMT-like phenotype, although the co-expression of inflammatory and immune-interacting genes, including the chemokines *CXCL1* and *CXCL8* and the NF-κB pathway component *NFKB1* suggests that inflammatory activation may also contribute to the observed profile.

Another cluster highly predominant in the RC-V ROs was the perivascular fibroblast cluster 2, characterised by high expression of fibroblast (*COL1A1, COL5A1, COLCA1, COLCA2, FN1, VCAN*) vascular and endothelial signalling markers (*VEGFA*, *ANGPT2*, *EGFL7*, *EMCN*, *NOS3*, *CXCL12*, *ANGPTL4*, *PDGFC*, *KLF2*, and *ECSCR*), alongside retinal vascular-associated fibroblast markers (*SOX18*) (**Table S1**). The transcriptomic profile of these cells points to associated mural cells involved in angiogenesis, vascular remodelling, and hypoxia responses. Notably, a distinct but closely located cluster of vascular smooth muscle cells (cluster 19) was also identified, further supporting the presence of complex vascular elements within the organoid structure.

We also identified a cluster characterised by high expression of canonical pericyte and mural-cell markers, including *PDGFRB, CSPG4* and *NOTCH3* (cluster 9, **Table S1**), together with a prominent contractile programme comprising *ACTA2, TAGLN, CNN1, MYL9, DES, MYOCD* and *MYOCD-AS1*. The cluster also expressed genes associated with vascular support and extracellular matrix (ECM) organisation, including *PDGFRA, COL4A1, COL4A5, COL4A6, LOX, SPARC, MFAP4, FBLN1, COL1A2* and *MRC2*, as well as regulators of vascular remodelling such as *TGFB2, TGFB3, ANGPTL2, VEGFD* and *ANGPTL4*. Collectively, this expression profile supports annotation of cluster 9 as pericytes/vascular mural cells. Notably, cluster 9 exhibited a pronounced ECM-remodelling and activated phenotype. Expression of *SERPINE1, MMP14, ADAMTS1, ADAMTS5, ADAMTS8, ADAMTS9, ADAMTS15, ADAMTS16, STC1, STC2, HMOX1, SOCS3* and *THBS1*, together with *CCN1, CCN3, FSTL1* and *SPARC* (**Table S1**), is consistent with enhanced matrix turnover and tissue-remodelling activity. The concurrent expression of *ACTA2, TAGLN, CNN1, MYL9* and *MYOCD* indicates a strongly contractile phenotype, while *TGFB2, TGFB3* and *TGFBR2* suggest engagement of TGF-β-associated signalling. In addition, expression of *CXCL1, CXCL3, CXCL6, IL33, IL4R, IFI30, IFITM1, BST2* and *ISG20* indicates an inflammatory and stress-responsive component to the transcriptional state. Together, these features are compatible with a contractile, ECM-producing and remodelling pericyte phenotype, rather than a quiescent vascular support state. Because this cluster was present across all experimental groups, it likely originated endogenously within the ROs rather than from the exogenously added vascular cells.

To infer potential cell–cell communication, LIANA consensus ligand–receptor analysis was performed between sender cell populations (perivascular fibroblasts-cluster 2 and activated inflammatory ECs-cluster 8) and recipient cell populations (vascular smooth muscle cells- cluster 19 and pericytes-cluster 9). Overall, both perivascular fibroblasts and activated inflammatory ECs participated in a dense communication network, sharing multiple signalling axes across both target cell types. However, activated inflammatory ECs (cluster 8) exhibited a broader and higher-magnitude interaction profile than perivascular fibroblasts (cluster 2), particularly toward pericytes (cluster 9), which served as the more communication-active recipient overall (**Fig. S8**). Prominent pathways identified across these interactions included pro-angiogenic an endothelial signalling (VEGFA paired with FLT1_KDR, CD44, EGFR, and EPHB2), lipid-handling and stress-response signalling (APOE interactions with LRP1, LSR, LRP5, and SCARB1), inflammatory crosstalk (MIF→ACKR3/EGFR and ICAM1→CAV1), and extracellular matrix remodelling (COL6A2 interactions with ITGA2_ITGB1, ITGA3_ITGB1, and CD44) (**Fig. S8**).

The RC-V ROs, which exhibited the highest predominance of activated inflammatory ECs and perivascular cells, were also characterised by a lower presence of all retinal cell types, corroborating our IF findings (**Fig. 7**). The combination of endothelial activation, perivascular mesenchymal remodelling, and retinal cell loss bears all the hallmarks of diabetic retinopathy (DR). To probe into this further, we compared our scRNA-seq clusters 2 and 8 to the transcriptomic dataset, GSE117238, from human retinal ECs (hRECs) treated with low glucose (LG), or high glucose (HG) generated by Shao and colleagues (Shao *et al*. 2019). To this end, we compared the differentially expressed genes in Cluster 2 and Cluster 8 with genes upregulated under HG conditions in the dataset reported by Shao *et al*. (2019). The HG gene set comprised 763 genes, of which 679 were unique to this comparison, while 103 genes were also represented among Cluster 8-associated genes and 84 among Cluster 2-associated genes (**Fig. S9A**).

Notably, the overlap was greater for Cluster 8 than for Cluster 2, with 103 genes shared between Cluster 8 and the Shao *et al*. high-glucose signature, compared with 84 genes shared between Cluster 2 and the high-glucose signature. A further 47 genes were common to all three gene sets, indicating a subset of genes consistently associated with the transcriptional response across the independent dataset and both clusters. In contrast, Cluster 2 and Cluster 8 shared a considerably larger set of genes (646 genes), suggesting that the two clusters retain substantial transcriptional similarity despite their differential relationship with the HG signature.

To determine whether these overlaps were greater than expected by chance, representation- factor analysis was performed (**Fig. S9B**). Cluster 8 showed significant enrichment for genes upregulated under HG conditions in Shao *et al*. (RF = 1.51, P = 1.60 × 10⁻⁵), indicating that these genes were represented within Cluster 8 at a frequency greater than expected by chance. In contrast, although Cluster 2 also showed an increased representation factor (RF = 1.32), this enrichment did not reach the predefined significance threshold of P < 0.001 (P = 0.00485). Thus, only Cluster 8 demonstrated significant enrichment for the HG-associated transcriptional programme, suggesting that this population more closely recapitulates aspects of the transcriptional response observed following HG exposure in Shao *et al*. (2019).

Together, these findings provide independent support for the interpretation of Cluster 8 as a population exhibiting a metabolic- and stress-responsive transcriptional state with features relevant to DR. The significant enrichment of the Cluster 8 transcriptome for genes induced by HG exposure in endogenously arising human retinal ECs suggests that this population captures aspects of the endothelial response to a diabetic-like metabolic environment. This is particularly notable given the prominent endothelial activation and perivascular remodelling observed in the RC-V ROs, phenotypic features that are consistent with the vascular dysfunction and tissue remodelling characteristic of DR. Although the relatively modest representation factor indicates that Cluster 8 does not recapitulate the complete HG transcriptional programme, the significant overlap across independent experimental systems supports the possibility that this population reflects molecular features of retinal vascular stress associated with diabetic pathology.

## Discussion

Here we show that VOrs can serve as a renewable source of endothelial, pericyte, astrocyte and microglia-like cells capable of migrating into developing ROs, and that transient integration of these cells, when timed appropriately, can be achieved without abolishing retinal lamination or light-evoked RGC output. At the same time, our data reveal an unexpected endogenous capacity of ROs to generate inflammatory endothelial-like and perivascular mesenchymal populations under pro-angiogenic culture conditions, yielding a transcriptional programme that partially recapitulates DR. Together, these findings delineate both the opportunities and the constraints of vascular–immune engineering in human ROs and highlight a previously underappreciated endogenous vascular–immune axis that emerges in standard long-term retinal cultures.

## Transient, early exposure to VOr-derived cells supports retinal development, but static niches limit vascular persistence

Our initial assembloid experiments confirmed that VOr-derived CD31⁺ ECs, GFAP⁺ astrocytes and IBA1⁺ microglia-like cells can exit the VOrs during VOr-RO fusion, invade the basal aspect of ROs and associate with nascent retinal tissue at day 90 of RO differentiation. However, retinal lamination deteriorated after ∼28 days, indicating that late-stage physical fusion is insufficient to generate a stable, laminated, vascularised retina. This is consistent with broader organoid literature showing that uncontrolled coalescence of distinct organoid types can disrupt tissue architecture and patterning (Kistemaker *et al*., 2025). By contrast, dissociating VOrs at defined timepoints (day 12 or day 30) and seeding them onto 2D retinal neuroepithelium prior to 3D transition allowed VOr-derived cells to be present transiently present until day 100 of differentiation. Under these conditions, particularly in the V30-RO group, we observed sustained lamination, correct anatomical positioning of major neuronal subtypes, and RGC light responses comparable to non-vascularised controls. These results suggest that early, transient exposure to VOr-derived cells, rather than persistent, high-density vascular co-culture, may be more compatible with normal retinal development *in vitro*.

The superior performance of V30-ROs relative to V12-ROs in functional assays may reflect a better match between the timing of vascular cell addition and the emergence of inner retinal circuitry. Day 30 VOrs contain a more mature complement of pericytes, smooth muscle-like cells and microglia, which could provide stabilising cues without overwhelming the developing neuroepithelium (Sun *et al*., 2022). In contrast, day 12 VOrs are enriched for proliferative endothelial progenitors that may exert stronger pro-angiogenic and remodelling pressures, potentially perturbing later synaptic maturation. This interpretation aligns with reports that excessive or mis-timed VEGF signalling can disrupt retinal lamination and RGC survival during retinogenesis (Hashimoto *et al*. 2006). In accordance, our scRNA-Seq data showed that RGCs were reduced in all the experimental groups that were supplemented with VEGF during the differentiation process.

Notably, VOr-derived cells seeded into the retinal neuroepithelium showed a sharp decline in survival within approximately 20 days post-addition. We attribute this primarily to an inappropriate niche and to central hypoxia in the organoid core, where diffusion-limited oxygen and nutrient supply are known to drive necrosis and apoptosis in inner cell layers. In larger, avascular organoids, cells beyond ∼200–400 µm from the surface become hypoxic or necrotic, a constraint that has been linked to progressive loss of inner-retinal neurons (including RGCs) and to destabilisation of any nascent vascular networks. Vascular ECs are particularly sensitive to hypoxia-induced apoptosis, so the absence of a perfused microcirculation likely accelerates their death once they are embedded deep within the organoid (Stempien-Otero *et al*., 1999).

A second, non-mutually exclusive explanation is the lack of physiological fluid shear stress and unidirectional flow, both of which are critical for EC survival, alignment, lumen formation, and vessel maturation. In static organoid cultures, the absence of convective flow prevents the establishment of stable shear-stress cues that normally promote endothelial gene expression, cytoskeletal remodelling, sprouting, and network persistence; without these cues, nascent vessels tend to regress or remain immature and leaky (Quintard *et al*., 2024). Microfluidic studies in kidney and brain organoids, as well as in vessel-on-chip systems, show that applying unidirectional laminar flow enhances endothelial coverage, vessel area, junctional density, and longevity, and that continuous perfusion is required to sustain sprouting and prevent retraction (Homan *et al*., 2019; Quintard *et al*., 2024). By contrast, static or disturbed-flow conditions favour endothelial dysfunction and vessel instability, which may explain why VOr-derived vascular cells fail to persist beyond the first month when embedded in a non-perfused ROs (Ryan *et al*. 2021).

Additional contributing factors likely include: (i) mismatched oxygen gradients, inner retina is physiologically hypoxic (∼2% O₂) while outer retina is more oxygenated (∼18% O₂) (Wangsa- Wirawan and Linsenmeier, 2003), a gradient that is difficult to reproduce in spherical, avascular organoids and that can impair inner-layer survival; (ii) accumulation of metabolic waste and limited nutrient exchange in the organoid core, exacerbating stress and cell death; and (iii) absence of axonal exit and appropriate neurovascular coupling, which *in vivo* supports RGC and inner-retinal maintenance but is lacking in closed organoid systems. Together, these niche deficiencies provide a mechanistic basis for the rapid attrition of VOr-derived cells after seeding into ROs.

## Endogenous mesodermal differentiation and the emergence of an inflammatory endothelial compartment

A striking and unanticipated finding was the appearance of CD31⁺ endothelial-like cells, perivascular fibroblasts, pericytes and αSMA⁺ smooth-muscle-like cells in ROs that never received exogenous vascular cells (RC-V group). This occurred in the context of a 1:1 retinal: endothelial media mix supplemented with VEGF, suggesting that pro-angiogenic conditions can drive endogenous mesodermal differentiation within hESC-derived retinal cultures. Single-cell RNA sequencing identified a prominent cluster (cluster 8) with an activated, inflammatory endothelial-like transcriptional profile, co-expressing residual endothelial genes (*CDH5, EMCN, ENG, NOS3, THBD*) and mesenchymal/extracellular matrix genes (*ACTA2, FN1, COL1A1, POSTN, MMPs, SNAI1*). This partial EndMT-like state, combined with chemokine and cytokine expression (*CXCL2, IL6, ICAM1*), mirrors features described in diabetic and hypoxic retinal endothelium.

Importantly, cluster 8 was significantly enriched for genes upregulated in human retinal ECs exposed to HG in an independent dataset, supporting the notion that our RC-V ROs spontaneously acquire a DR-like endothelial stress signature in the absence of exogenous glucose manipulation. However, the elevated glucose levels (∼25mM) in retinal media used from day 84 of differentiation onwards, may have contributed to the DR-like transcriptional phenotype observed in RC-V ROs, consistent with the findings of a recent study by Bertelli *et al*., (2022) demonstrating that sustained glucose levels (∼25mM) induce premature senescence and functional impairment in human retinal ECs, including reduced tubulogenic and barrier capacity. This finding has two implications. Firstly, it cautions that pro-angiogenic media formulations, while useful for promoting vascular integration, can inadvertently generate pathological endothelial states that confound disease modelling or therapeutic testing. Secondly, it reveals that human ROs possess an intrinsic capacity to generate vascular-like and perivascular mesenchymal populations when provided with appropriate cues, challenging the assumption that all vascular elements must be exogenously supplied.

Notably, activated inflammatory endothelial-like cells and perivascular fibroblasts were absent from ROs seeded with VOr cells, despite the identical culture conditions and VEGF supplementation. In contrast, RC-V ROs exhibited a marked reduction in all major retinal cell lineages, accompanied by disrupted retinal lamination. These abnormalities were not observed in ROs seeded with VOr cells, suggesting that the incorporation of exogenous vascular cells may support retinal development while also preventing the inflammatory activation of endogenous endothelial-like cells within the ROs. These findings have important implications for the future development of vascularised ROs, highlighting the potential benefit of combining exogenous vascular cells with components that promote a supportive pro-endothelial niche to achieve more physiologically relevant vascularisation.

## CD31 is detectable on ATOH7^+^ neurogenic progenitors during early retinal organoid development

To determine whether endothelial marker expression overlaps with early neurogenic programmes in human ROs, we co-stained developing organoids for the pan-endothelial adhesion molecule CD31 and the proneural transcription factor ATOH7, which marks the T1 transient neurogenic progenitors (Dorgau *et al*. 2024). From approximately day 30 to day 90 of differentiation, we observed a population of cells that were positive for both ATOH7 and CD31. These double-positive cells were predominantly located in the outer neuroblastic-like zone of the organoids, consistent with the spatial distribution of exiting progenitors and early neuroblasts described in single-cell transcriptomic maps of fetal retina and organoids. Within this ATOH7^+^ compartment, CD31 immunoreactivity was typically weaker and more punctate than the strong, continuous membrane staining seen in CD31hi endothelial tubes, suggesting a distinct, low-level expression state rather than full endothelial identity.

This pattern aligns with evidence from the developing brain that neural progenitors can express “extra-neural” markers, including CD31, CD34, and Tie2, in addition to classical neural markers such as Nestin. In embryonic mouse cortex, CD31+ cells associated with vessels co-express Nestin, and sorted CD31+ fetal cortical cells can form neurosphere-like clusters and differentiate into neural lineages in vitro, indicating bidirectional plasticity between vascular and neural compartments during development (Parati *et al*, 2002). Moreover, embryonic stem cell derivatives include a discrete Nrp1+/Flk1+ (VEGFR2+) population that can adopt either endothelial or neural fates depending on microenvironmental cues, supporting a developmental programme in which early progenitors carry both angiogenic and neurogenic potential (Gualandris *et al*, 2009). Tight functional coupling between CD31+ ECs and Nestin^+^ neural progenitors in neurogenic niches further reinforces the interweaving of progenitor and endothelial programmes during CNS development.

In the retina, our own work has shown that a subset of ATOH7+ neurogenic progenitors expresses low-level CD31 during the window of RGC genesis (Dorgau *et al*, 2024). This finding is consistent with the established role of ATOH7 in guiding progenitor competence and promoting RGC commitment while simultaneously regulating pathways that intersect with vascular development (Atac *et al*. 2024); ATOH7 loss-of-function not only abolishes RGCs but also leads to aberrant retinal vessel patterning, indicating that ATOH7+ progenitors are embedded in angiogenic programmes (Edwards *et al*., 2012). ATOH7-expressing retinal progenitor cells are highly responsive to VEGF, which modulates their proliferation, cell-cycle progression, and neuronal differentiation, and the vascular niche further shapes their survival and fate choices. CD31 is a core component of endothelial mechanotransduction and angiogenic signalling complexes that include VEGFR2, and it contributes to endothelial migration, tube formation, and VEGF-driven angiogenesis. In this context, transient, low-level CD31 expression on angiogenically “primed” ATOH7+ neurogenic progenitors is biologically plausible and may reflect a developmental state in which neurogenic and vascular signalling programmes are still intertwined before full fate restriction. Together, these data indicate that a subset of ATOH7^+^ neurogenic progenitors in early ROs exhibits low-level CD31 expression, consistent with a transient neuro-angiogenic state during the period when neurogenesis and vasculogenesis are still intertwined.

## Endogenous microglia-like and BAM populations

IBA1⁺ cells were present in all experimental groups from relatively early stages through to Day 210, with morphologies ranging from ameboid to ramified and expression of microglial markers (P2RY12, TMEM119, TREM2) at specific timepoints. Because labelled VOr cells were not used in our initial experiments, we cannot definitively determine whether these microglia-like cells and BAMs arose endogenously or were derived from the VOrs. However, their presence across all experimental groups suggests that they are most likely endogenous. Notably, we could not identify these populations by scRNA-seq, indicating that they are either present at very low abundance or preferentially lost during the single-cell dissociation procedure, consistent with our previous findings in ROs supplemented with exogenous microglia (Chichagova *et al*. 2023).

Partial co-localisation with the BAM marker CD163 further suggests that our ROs harbour a heterogeneous myeloid compartment comprising both parenchymal microglia-like cells and BAM-like populations. This aligns with recent work showing that microglia-like cells can arise intrinsically in unguided RO differentiation, localising to cystic, mesenchymal-like compartments adjacent to the neuroepithelium (Bartalska *et al*., 2022). The temporal dynamics we observe - peaking reactive morphologies around day 120, followed by more ramified, homeostatic-like profiles by day 180 in RC-ROs- are consistent with a developmental wave of synaptic pruning and circuit refinement, followed by stabilisation.

The presence of endogenous microglia-like and BAM-like cells has important consequences for experimental design. On one hand, it provides an opportunity to study neuroimmune interactions without exogenous cell addition. On the other hand, it complicates interpretations of studies that assume ROs are immune-naïve unless microglia are explicitly added. Our data suggest that future work should routinely profile myeloid populations in ROs, even in ‘non-immune’ conditions, and consider how culture conditions (e.g., BMP4 exposure, cyst formation, matrix composition) influence the balance between microglial and BAM-like states.

## Implications for modelling DR and other vascular retinopathies

The convergence of inflammatory endothelial activation, perivascular fibroblast expansion, ECM remodelling and retinal cell loss in RC-V ROs recapitulates several hallmarks of early DR, including endothelial dysfunction, neuroinflammation and glial reactivity. Unlike hyperglycaemia-based DR models that rely on exogenous glucose elevation, our system appears to generate a DR-like endothelial state through endogenous differentiation under pro-angiogenic conditions. This could be leveraged as a complementary model of metabolic–vascular stress, particularly for dissecting cell-intrinsic endothelial responses and their crosstalk with pericytes, smooth muscle-like cells and microglia.

However, several caveats apply. Firstly, our ROs still lack a perfused, lumenised inner retinal capillary bed with a defined inner BRB; the endothelial-like cells we observe are not organised into a physiological vascular plexus. Furthermore, the DR-like signature is partial and may not capture later-stage features such as neovascularisation, acellular capillaries or clinically relevant haemodynamic changes. In addition, the absence of systemic metabolic inputs (e.g., circulating lipids, immune cells, glycation end-products) limits the extent to which our model can replicate the full spectrum of human DR. Nonetheless, the significant transcriptional overlap with high-glucose-treated human retinal ECs supports the utility of this system for probing early endothelial stress pathways and testing interventions that target endothelial activation, EndMT or neuroimmune crosstalk.

## Limitations and future directions

Several limitations of our current approach should be acknowledged. Although VOr-derived cells can integrate into ROs, their long-term persistence is limited, with only a minority of organoids retaining detectable vascular cells by day 210. This likely reflects the absence of perfusion, physiological shear stress and stable basement membrane cues that are required for mature vessel maintenance. Strategies such as microfluidic perfusion, *in vivo* transplantation, or engineered extracellular matrices may be needed to stabilise these networks over longer periods. Secondly, our scRNA-seq dataset, while informative, is based on a modest number of cells and organoids; larger-scale, multi-timepoint atlases will be required to fully resolve the trajectories of endogenous endothelial-like and perivascular populations. Finally, we did not perform lineage tracing to definitively distinguish endogenous versus exogenous origins of specific vascular and immune subsets within V12- and V30-ROs; future studies using cell labelling, genetic barcoding or CRISPR-based lineage recording will be essential to map these fates with precision.

Looking forward, several avenues appear promising. Combining timed VOr dissociation with microfluidic or organ-on-chip platforms could provide both early vascular–immune exposure and sustained perfusion, potentially yielding stable, laminated, vascularised ROs. Targeted modulation of VEGF, TGF-β and inflammatory signalling during the window of vascular integration may help to suppress pathological EndMT-like states while preserving beneficial pro-survival cues. Finally, integrating metabolic perturbations (e.g., fluctuating glucose, advanced glycation end-products) with our endogenous inflammatory endothelial model could generate a more comprehensive *in vitro* platform for early DR, enabling high-content screening of neuroprotective and vasoprotective therapies.

## Concluding remarks

In summary, our work demonstrates that VOrs can be harnessed to transiently seed developing ROs with diverse vascular and immune cell types, and that careful timing of this exposure can preserve retinal architecture and function. At the same time, we uncover an endogenous, pro-inflammatory endothelial and perivascular programme that emerges under pro-angiogenic culture conditions and partially mirrors DR. These insights refine our understanding of what is required, and what must be restrained, to build physiologically faithful, vascularised and immunocompetent human retinal models for development, disease and therapy.

## Methods

### hESC and hiPSC culture

hESCs were maintained at standard cell culture incubator conditions at 95% humidity, 5% C02 and 37°C. H9 CRX-GFP and H9 hESCs to establish ROs and VOrs, respectively. H9 hESCs were purchased from WiCell Inc. with an MTA renewal approved annually for their intended use, whilst H9 CRX-GFP hESCs were generated in our team (Collin *et al*. 2016). For the live cell tracking experiment, H9 CRX-GFP and WT2 mCherry hiPSCs were used for the generation of ROs and VOrs, respectively. WT2 hiPSCs were generated in our team (Buskin et al. 20018) and further modified by lentiviral transduction to generate mCherry WT2 hiPSCs. hESCs/hiPSCs were grown on Matrigel™ Growth Factor Reduced Basement Membrane Matrix (Corning) using mTeSR™ 1 cell culture media (STEMCELL Technologies) supplemented with 1% penicillin-streptomycin (Thermo Fisher Scientific). Cells were passaged at a 1:3 or 1:6 ratio using 0.02% Versene-EDTA (Lonza) after a 3-minute incubation at 37 °C. Media was replaced daily. All nascent cultures were routinely tested for mycoplasma using the MycoAlert® Mycoplasma Detection Kit (Lonza, LT07- 118).

### VOr differentiation

To induce VOr formation, we followed the protocol of Sun et al, 2022. In brief, hESCs cultured in 6 well-plates were washed with PBS, dissociated using Accutase at 90% confluence (Gibco, Thermo Fisher Scientific) and seeded at a density of 9000 cells/well on V-bottom Lipidure pre-coated 96-well plates (Helena Biosciences) in the presence of 10 μM ROCK inhibitor (Y27632, Tocris). Upon embryoid body formation (D2), mTeSR™ 1 was completely aspirated, and spheroids were fed with APEL2 medium (STEMCELL Technologies) supplemented with 6μM CHIR99021 (Selleck) to promote mesoderm specification. On Day 4, endothelial differentiation was induced by the addition of 50 ng/ml VEGF (STEMCELL), 25 ng/ml BMP4 (STEMCELL) and 10 ng/ml bFGF (STEMCELL) to APEL2 media (STEMCELL). From Day 7, developing VOrs were maintained in MV2 medium (PromoCell) augmented with 50 ng/mL VEGF (STEMCELL Technologies), with media changes every other day until Day 12. At Day 12, VOrs were embedded in 20 μL Matrigel™ droplets (Corning) on parafilm-lined 1 mL pipette tip rack scaffolds (2-hour incubation at 37°C), and VEGF concentration in VOr expansion medium was reduced to 20 ng/mL (**Table S1**). From Day 16 to Day 30 the VOrs were placed on a shaker platform at 30rpm in the incubator, and VOr expansion media was substituted with VOr maturation media until D40 of VOr differentiation (**Table S1**).

### RO Differentiation

hESCs were differentiated to ROs using the 2D-3D method outlined by West *et al*. protocol (2022). In brief, upon reaching 90% confluence, hESCs were cultured in TeSR™-E6 for 2 days to prime retinal fate (Day 0-Day 2). From Day 3–Day 44, 2D retinal epithelium sheets were maintained in proneural induction medium (PIM) (Advanced DMEM/F-12 + 1% NEAA, 100 μM glutamine, 1% N2, 1% Pen/Strep) until formation of translucent horseshoe-shaped patches was evident (week 3-5). Day 12 and day 30 VOrs were dissociated using 25 U/mL papain (STEMCELL) and plated as single cells (200,000 cells/well) onto the 2D sheets at day 30 of retinal differentiation. Epithelium 2D patches were scratched in a grid-like pattern at Day 44 and transferred to ultra-low-attachment 6-well plates (Thermo Fisher Scientific). The control organoids (RCs) were maintained in RDMII medium until Day 70 (DMEM/F12, 2% B27 wo vit A, 10% FBS, 0.1 mM taurine, 2 mM GlutaMAX™,1% Pen/Strep), and RDMIII from Day 71 to Day 84 (RDM II + 1 μM fresh retinoic acid). From Day 84 until Day 220 RC-ROs were cultured in ALT medium (Advanced DMEM/F-12; 1% Pen/Strep, 2% B27-minus A, 1% N2, 0.1 mM taurine, 4 mM GlutaMAX™, 7.5 mM D-glucose, 12.5 μM non-fatty acid BSA, 1 μM RA withdrawn after Day 119). A second group of adapted retinal controls (RC-V ROs) and vascularized retinal organoids (V12- ROs, V30-ROs) were maintained in 1:1 retinal medium:MV2 (PromoCell) supplemented with 20ng/mL VEGF (STEMCELL Technologies) from the point of scratching until day 44, whereby the VEGF concentration was reduced to 10ng/mL until the end of differentiation (Day 220) (**Figure S2B**). Partial media changes were carried out 3x a week over the course of retinal differentiation.

### Cryosectioning of Retinal and Vascular Organoids

Organoids were collected at specified timepoints (days 60, 90, 120, 180 and 210 for ROs and day 40 for VOrs), fixed in 4% paraformaldehyde (PFA)/5% sucrose in PBS for 20 minutes at 4°C. For VOrs special care was taken for transfer using spoons appropriated from Pasteur pipettes. Organoids were dehydrated at 4°C in a successive gradient of sucrose (6.25%; 12.5% and 25% overnight) and embedded in OCT (CellPath; 5 organoids per block) before snap-freezing on dry ice. Vascularised retinal organoids/controls and VOrs were cryosectioned at 10-12 µm and 20-30 µm, respectively (Leica CM1950), thaw-mounted onto Superfrost Plus™ slides (Epredia™), air-dried 1 hour at RT, and stored at −20°C before further processing.

### Immunohistochemistry

Cryosections of vascularised ROs, ROs or VOrs were briefly rehydrated in PBS (3 × 5 minutes) and blocked in 10% donkey or goat serum in 0.3% Triton X-100/PBS for 1h at RT. Primary antibodies (**Table S1**) were diluted in antibody buffer (0.3% v/v Triton X-100, 1% w/v BSA in PBS) and incubated overnight at 4°C in a humidified chamber. The next day, slides were washed with antibody buffer (3 × 10 minutes), incubated with species-appropriate Alexa Fluor® secondary antibodies (1:800 for 546 and 1:500 for 647 Alexa Fluor® antibodies, Thermo Fisher Scientific, **Table S1**) for 1 hour at RT. Sections were washed again (3x15 minutes), and mounted in VECTASHIELD® H-1000 (Vector Laboratories) containing Hoechst (1:1000; Thermo Fisher Scientific). Fluorescence imaging was enabled by optical sectioning on an Axio Imager microscope (Zeiss, Apotome, 10-15 Z-stacks). Secondary controls omitted primary antibodies.

### Tissue clearance and confocal imaging

Wholemount vascularised retinal and vascular organoids were cleared following the Visikol® HISTO-M™ (Visikol Inc.) protocol. Organoids were fixed in 4% PFA for 30 minutes and washed twice in PBS (15 minutes). The samples were transferred to imaging vessels and permeabilised via DMSO/methanol dehydration to prepare for antigen retrieval (20% DMSO/methanol [10 minutes], 80% methanol [10 minutes], 50% methanol/PBS [10 minutes], 100% PBS [10 minutes], 0.2% Triton X-100/PBS [10 minutes]). Once permeabilised, organoids were treated with Visikol® HISTO™ Penetration Buffer (Visikol Inc., 37 °C, 30 minutes) and subsequently blocked (Visikol® HISTO™ Blocking Buffer, Visikol Inc.) under gentle shaking at 37 °C for 1 hour. Incubation with primary antibodies (Visikol® HISTO™ Antibody Buffer (**Table S1**) was carried out on a shaker platform for 1 hour at 37 °C, followed by 3x10 minute washes Visikol® HISTO M™ Washing Buffer). Secondary antibodies and Hoechst nuclear stain (1:1000) were applied simultaneously in Visikol® HISTO™ Antibody Buffer for 1 hour at 37°C. This was followed by 5x5 minute washes (Visikol® HISTO™ Washing Buffer, Visikol inc.) and final pre-clearing dehydration steps with methanol (50%, 80%, 100%; 10 minutes each). Wholemounts were ultimately cleared for 15 minutes at RT (Visikol® HISTO-M™, Visikol Inc.) and imaged at confocal resolution on a Zeiss LSM 880 Airyscan microscope using 20×/10x objectives (Z-stacks at 1 µm intervals). Secondary only controls were included to confirm the specificity of primary antibodies.

### Flow cytometry

Papain stock (250 U/mL) was activated at 37°C for 30 minutes (ddH₂O, 5 mM L-cysteine-HCl, 1.1 mM EDTA, 2.44 µL β-mercaptoethanol). ROs and VOrs were washed in HBSS, dissociated with papain (250 U/mL and 25 U/mL HBSS, 250 µl DNase I for ROs and VOrs, respectively) and neutralised with inhibitor solution (20% FBS in HBSS). Dissociated cells were strained (100 µm) and counted using Countess™ II FL Automated Cell Counter (Invitrogen, Thermo Fisher Scientific, Countess™ cell counting chamber slides, Thermo Fisher Scientific). VOr samples were FcR-blocked (20% Blocking reagent, 10% FBS HBSS flow buffer) for 10 minutes (4°C) and stained with antibodies in flow buffer for 30 minutes (4°C), avoiding light exposure (**Table S1**). Finally, samples were washed (flow buffer 3×1100 rpm, 5 minutes, 4°C) and resuspended in 500 µl flow buffer supplemented with Hoechst (1:1000) for nuclei counterstaining. Flow cytometry analysis was conducted on the BD LSRFortessa (BD Biosciences) using unstained and isotype controls. At least 10,000 events were collected for each sample.

### qRT-PCR

Total RNA was extracted following the protocol for an SV Total RNA Isolation System kit (Promega, Cat. No. Z3101). RNA concentrations were quantified by NanoDrop™ microvolume spectrophotometer (Thermo Fisher Scientific), and 1 µg aliquots were reverse-transcribed to cDNA using a GoTaq® 2-Step RT-qPCR System kit (Promega). Equal amounts of cDNA template (1:10 dilution) were used to set up a 384-well qPCR reaction using a PowerTrack™ SYBR® Green master mix (Applied Biosystems) with target-specific primers in a 384-well format on a QuantstudioTM 7 Flex Real-Time PCR system (Applied Biosystems). Data were normalised to the expression of *GAPDH,* and gene expression was calculated using the 2^(-ΔΔCT) method. Samples were run in triplicates and data were presented as means + SEM. Primer oligonucleotide sequences are shown in **Table S1**.

### scRNA-Seq

Prior to processing, Papain (STEMCELL Technologies 7466) was reconstituted using an activation solution containing 0.067 mM mercaptoethanol (Gibco 21985-023), 1.1 mM EDTA (Invitrogen 15575020), and 5.5 mM L-cysteine HCl (Thermo Fisher Scientific, L06328.14) and filter-sterilised. Papain was diluted to a stock concentration of 250 units/mL and activated at 37°C for 30 minutes. Immediately before organoid dissociation, a dissociation solution was prepared by diluting 250 unit/mL papain stock solution to a value of 30 units/mL with 125 units/mL DNase I (STEMCELL Technologies, #07900) in HBSS (STEMCELL Technologies, #37150). Up to a maximum of 10-day, 210-aged organoids were harvested and washed once using HBSS. Following this, organoids were suspended in 500 μL dissociation solution and incubated at 37°C using an orbital shaker at 90 rpm or higher for 30 minutes. Following 30-minute incubation, organoids were manually triturated using a pipette 10 times, taking care to avoid introduction of bubbles. If dissociation was insufficient, samples were returned for further incubation. Following complete dissociation, the organoid suspensions were passed through a 37 μm reversible cell strainer (STEMCELL Technologies, #27215) directly into an inhibitor solution composed of 20% FBS in HBSS. For each sample, 20,000 single cells were loaded onto a Chromium Next GEM Chip G (10x Genomics, 1000120) and encapsulated using the Chromium Next GEM Single Cell 3′ Reagent Kit (v3.1, 10x Genomics, 1000268) following the manufacturer’s protocol. Sequencing libraries were constructed using the Chromium Single Cell 3′ Library C Gel Bead Kit (v3.1, 10x Genomics, 1000121). The prepared scRNA-seq libraries were subsequently sequenced on an Illumina NovaSeq 6000 platform to achieve a target depth of 50,000 reads per cell.

### scRNA-Seq analyses

BCL files were demultiplexed into sample FASTQ files using bcl2fastq2 (v.2.20.0). Transcripts were then quantified using 10x Genomics Cell Ranger (v.9.0.1, (Zheng *et al*. 2017)) via alignment to GRCh38 (GRCh38-2024-A supplied by 10X Genomics).

Output files from the Cell Ranger pipeline were imported into R (R Core Team 2025) using Seurat (v.5.3.1) and quality controlled following Dorgau *et al*. (2024). Briefly, cells that had fewer than 1,000 reads or 500 genes and more than 10% mitochondrial reads were removed. Probable doublets were identified and removed using DoubletFinder (v2.0.6, (McGinnis *et al*. 2019)). Each sample was quality controlled independently following the standard clustering workflow of Seurat. During the scaling process the following factors were regressed out "percent.mt", "nCount_RNA", "nFeature_RNA".

Samples were then integrated and batch corrected using Harmony (v.1.2.4). Following integration, differentially expressed cluster markers were generated using the FindMarkers function from Seurat to allow for cell type annotation. Consensus ligand–receptor analysis was completed using Liana (v0.1.14 (Dimitrov *et al*. 2024)). Further downstream visualisation was completed using SeuratExtend (v.1.2.8 (Hua *et al*. 2025)).

### RNA-seq Shao et al (GSE117238)

FASTQ files from GSE117238 were downloaded from the SRA and analysed using nf-core/rnaseq (v3.22.2). Merged transcript count data was then imported into R and differentially expressed genes identified using DESeq2 (v1.52 (Love *et al*. 2014)) and compared to the scRNA-seq data. Following Kim *et al*. (2001) a representation-factor analysis was then performed in R to compare the overlap of genes between Shao *et al*. and cluster08 and cluster02 (Kim *et al*. 2001; Patraquim *et al*. 2022).

### MEA

Multielectrode array (MEA) recordings were performed on the BioCamX platform using the high- density MEA CorePlate™ 1W 27/42L (3Brain GmbH, Lanquart, Switzerland), which integrates 4096 square microelectrodes in a 2.67 × 2.67 mm area and 42 μM inter-microelectrode spacing. 24 hours prior to MEA recordings, 9-cis retinal (10 nM; Sigma-Aldrich, UK) was added to the incubation medium. ROs were transferred to 33 °C artificial cerebrospinal fluid (aCSF) containing the following (in mM): 118 NaCl, 25 NaHCO3, 1 NaH2PO4, 3 KCl, 1 MgCl2, 2 CaCl2, 10 glucose, 0.5 L-Glutamine, and 0.01 9-cis-retinal. Organoids were opened longitudinally and placed with the presumed retinal ganglion cell (RGC) layer facing down onto a 4096-channel multielectrode array (MEA 3Brain GmbH, Lanquart, Switzerland). They were flattened using a translucent polyester membrane filter (Sterlitech Corp., Kent, WA, USA) with a stainless-steel ring of 3 mm diameter on top. The organoids were allowed to settle for at least 2 hours while continuously perfused (1 mL/min) with bubbled aCSF. Recordings were filtered with a 50 Hz high-pass filter in BrainWaveX (3Brain) and stored in an HDF5 format. Action potentials (spikes) from retinal organoid ganglion cells were detected and sorted using Herdingspikes2 (https://github.com/mhhennig/HS2) as described in Hilgen *et al*. (2017). Briefly, spikes were first identified as threshold crossings on each channel, then merged into unique events based on spatial and temporal proximity. For each detected spike, a location was estimated from the signal’s centre of mass. Spike sorting involved clustering all events using a feature vector that includes the locations and the first two principal components of the largest waveform.

Light stimuli were projected onto the retina as described elsewhere (Hilgen *et al*., 2022). Briefly, light stimuli were projected using a DLP LightCrafter (Texas Instruments, TX, USA) and custom optics onto the 2.67 x 2.67 mm field of the MEA chip, where the retinal organoids are placed. A full-field stimulus with increasing contrasts (1-second contrast steps, max irradiance 217 μW/cm2, 1 second black, followed by 1 second 33% grey, 1 second black, 1 second 66% grey, 1 second black, and 1 second white) was used to detect potential basic light-driven responses in the retinal organoids. cGMP (cyclic guanosine monophosphate) acts as a second messenger in photoreceptor cells, regulating the phototransduction cascade. In darkness, high cGMP levels keep cyclic nucleotide-gated (CNG) channels open, allowing steady depolarising inward current. cGMP (8- bromoguanosine 3′,5′-cyclic monophosphate, Sigma-Aldrich, MO) was puffed in the recording chamber (final concentration: 100 μM), and activity was recorded continuously 2 min before the puff and after. GABA functions as an inhibitory neurotransmitter in the retina, playing a crucial role in regulating information flow and influencing the retinal response to light. When GABA is applied, it activates GABA receptors, altering the physiological properties of retinal organoid cells. GABA (γ-aminobutyric acid, Tocris Bioscience, Bristol, UK) was puffed in the recording chamber (final concentration: 125 μM), and activity was recorded continuously 2 min before the puff and after. Statistical significance and firing rate analyses were evaluated by using MATLAB (Mathworks, MA, USA) and Prism (GraphPad, CA, USA). RGCs were considered responsive if they showed at least a 25% increase or decrease in spiking activity during a 90 s time window during stimulus (light, cGMP, GABA) compared to a similar time window before the stimulus (spontaneous activity). For each cell, all spikes occurring during these time windows were counted, and the mean percentage change (± SD) in activity between windows was calculated.

### Generation of mCherry WT2 hiPSCs

*E. coli* containing the EF1A>mCherry/Puro construct (VectorBuilder; VB900200-4194vat) were cultured in LB and plasmid DNA purified by miniprep (ThermoScientific, K0503) after 24 hours. The mCherry, packaging (PAX2) and envelope (VSV-G) plasmids were co-transfected into HEK293T cells at ∼80% confluency and incubated overnight. The following day, medium was replaced with 293T medium (MEM + 10% FBS, 2 mM L-glutamine, 1% Pen/Strep), and supernatants containing lentiviral particles were collected at 48 and 72 hours, then concentrated with PEG solution (Merck, P2139). Concentrated mCherry lentivirus was aliquoted and stored at −80 °C. WT2 hiPSCs were transduced at ∼30% confluency, and 72 hours later mCherry-positive cells were selected with puromycin (0.5 µg/ml) to obtain a homogeneous mCherry+ population (Buskin *et al*., 2018). Cells were imaged on an Axio Observer Z1 inverted fluorescence microscope to confirm all cells were positive, and at ∼80% confluency were frozen and stored in liquid nitrogen.

## Declarations

### Ethics approval and consent to participate declaration

The hESC H9 (WA09) was purchased from WiCell Research Institute (Madison, WI, USA). WiCell has confirmed that the original derivation of H9 was performed under ethical approval and that informed consent was obtained from the embryo donors; the line is NIH-approved (NIHhESC-10-0062).

Human fibroblasts (WT2) were purchased from Lonza (Basel, Switzerland) and reprogrammed into WT2 hiPSCs in our laboratory (Buskin *et al*. 2018). Lonza states that all human primary cells are ethically sourced, with tissue collection conducted under IRB/ethics-approved protocols and documented informed consent from donors, in compliance with applicable regulations.

### AI Declaration

AI was used solely for grammar checks during the writing of this article.

## Author contributions

RT- experimental design, data collection and analyses, figure preparation, manuscript writing RS, JK, VB, BD, RH, JC, SW – data collection and analyses

MDT – bioinformatic data analyses, figure preparation

GH – data collection and analyses, figure preparation, fundraising RZ, RM, LA – study design, fundraising, dissemination

ML – study and experimental design, data analyses, figure preparation, manuscript writing, fundraising, dissemination

All authors approved the final version of the manuscript.

## Data availability

Single cell RNA-Seq data were submitted to the Gene Expression Omnibus under the accession number **GSE346651.** To review GEO accession GSE346651: Go to https://eur03.safelinks.protection.outlook.com/?url=https%3A%2F%2Fwww.ncbi.nlm.nih.gov%2Fgeo%2Fquery%2Facc.cgi%3Facc%3DGSE346651Cdata=05%7C02%7Cmajlinda.lako%40newcastle.ac.uk%7C1da910f930ba48121fba08df0f154302%7C9c5012c9b61644c2a91766814fbe3e87%7C1%7C0%7C639246256162068059%7CUnknown%7CTWFpbGZsb3d8eyJFbXB0eU1hcGkiOnRydWUsIlYiOiIwLjAuMDAwMCIsIlAiOiJXaW4zMiIsIkFOIjoiTWFpbCIsIldUIjoyfǪ%3D%3D%7C0%7C%7C%7CCsdata=tvcPPAWje3I1ltGXlAGEbNlSbrvaf0GaxoRqAhHq1cw%3DCreserved=0 Enter token opgvuieajzenved into the box

