## Supplementary material for "Transient Vascular Cell Integration and Intrinsic Media-Stimulated Endothelial Activation Reveal Diabetic Retinopathy Signatures in Human Retinal Organoids": Suppl Info

Prof. Majlinda Lako

Biosciences Institute,

Newcastle University,

Newcastle NE1 3BZ

United Kingdom

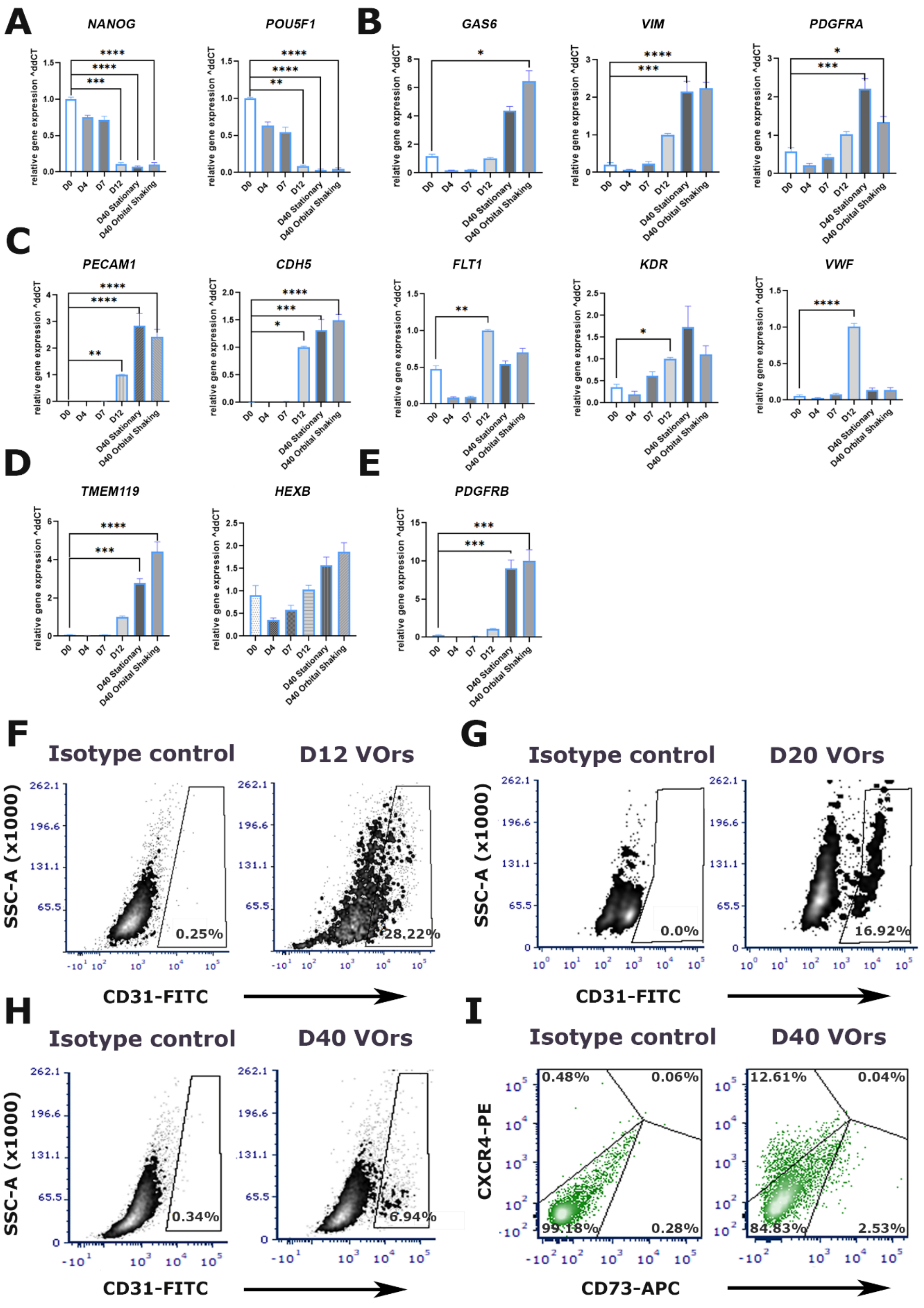


**Fig. S1: Longitudinal assessment of VOr differentiation by qPCR and flow cytometry. A.** qPCR analysis indicates successful VOr differentiation as evidenced by the reduction in *NANOG* and *POUF5F1* gene expression in D40 VOrs. **B.** Conversely, a significant increase in the expression of astrocytic marker *GAS6*, *VIM* and *PDGFRA* is seen in D40 VOrs. **C.** qPCR panel corroborates endothelial cell differentiation as shown by the gradual increase in endothelial markers *PECAM1*, *CDH5*, *KDR* and a peak of *FLT1* and *VWF* by D12 of vascular development. **D.** qPCR shows an increase in microglial marker *TMEM119* expression in D40 VOrs cultured under both stationary conditions and mechanical agitation relative to D0 VOrs. No significant difference between *the* stationary conditions and mechanical agitation is observed for the microglial marker *HEXB* expression. **E.** Pericyte gene marker *PDGFRB* is significantly upregulated in mature D40 VOrs by qPCR relative to day 0 samples. Data were analysed by one-way ANOVA with Tukey's multiple-comparisons test. Data are presented as mean ± SEM; n=3 differentiations. **F-G.** Forward and side scatter plots on flow cytometry show a peak of CD31-FITC+ expression in D12 VOrs (28.22%), followed by a gradual drop in D20 VOrs (16.92%) and mature D40 VOrs (6.94%) relative to the isotype control. **I.** Flow cytometry analysis of day-40 (D40) VOrs cultured under mechanical agitation revealed a greater proportion of CXCR4–PE$^+$ arterial-associated cells (12.61%) than CD73–APC$^+$ venous endothelial cells (2.53%). Data represent mean ± SEM (n = 3, ***p < 0.001, one-way ANOVA).

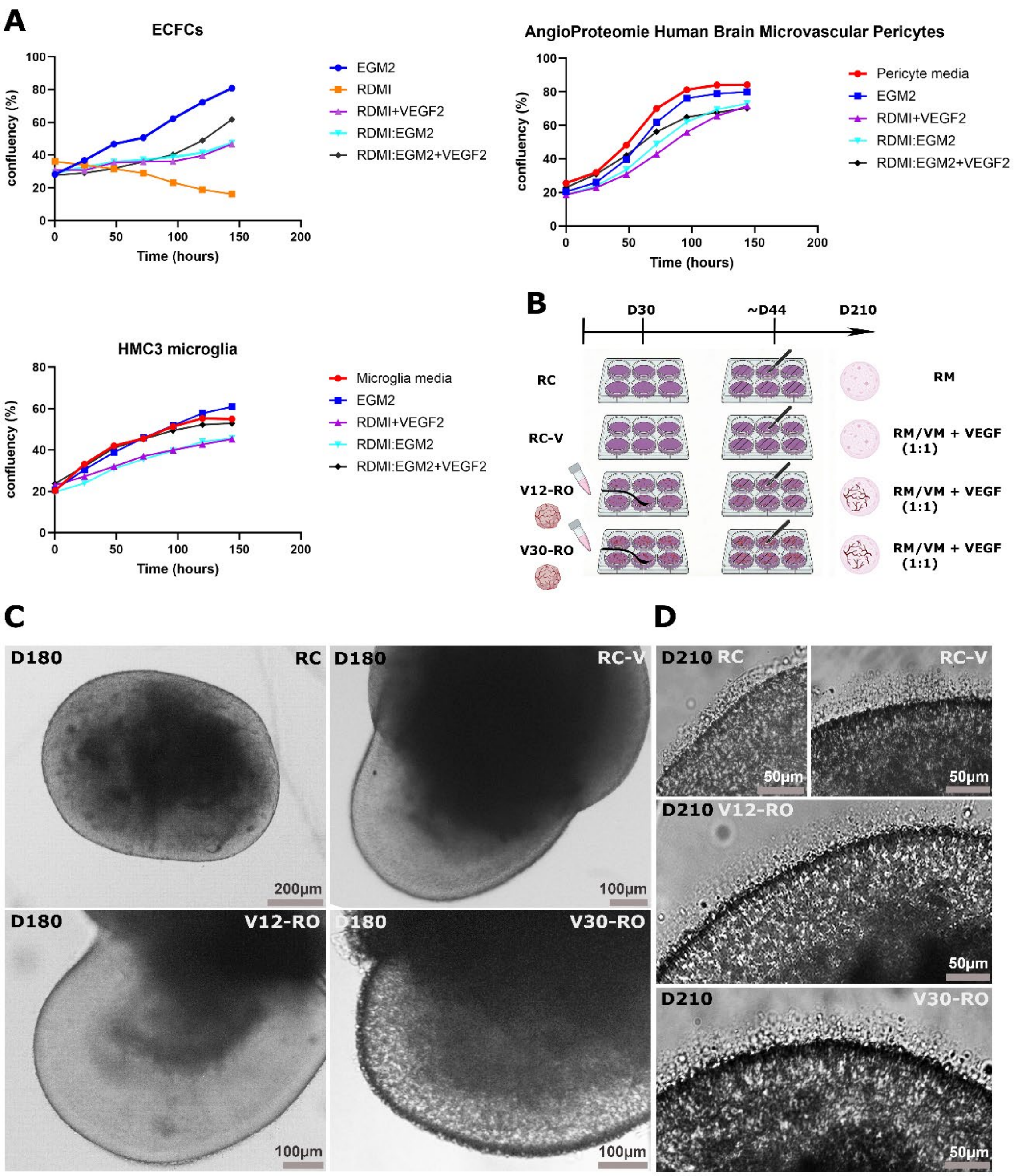


**Fig. S2: RDMI supplemented with EGM-2 and VEGF supports the highest viability of endothelial colony-forming cells (ECFCs). A.** Confluency plots showed comparable ECFC survival in EGM-2 endothelial medium and in a 1:1 RDMI: EGM-2 medium supplemented with VEGF over the course of a week. Angio-Proteomie human brain microvascular pericytes and HMC3 microglia displayed comparable performance across the endothelial-tailored culture media tested. Results are presented as percentages (%). **B.** The schematic outlines a four-arm experimental design comprising control RC ROs, media-control RC-V ROs, V12 ROs, and V30 ROs. Media-control RC-V ROs were cultured in a 1:1 retinal medium: vascular medium (RM: VM) mixture supplemented with VEGF from D30 until the end of retinal differentiation. To generate V12 and V30 ROs, VOrs aged D12 or D30, respectively, were dissociated and added to the developing retinal neuroepithelium at D30 of RO differentiation. The resulting structures were lifted at D44 to establish three-dimensional ROs and cultured in RM: VM (1:1) mix supplemented with VEGF until the end of differentiation. **C.** Representative brightfield images of mature D180 RC, RC-V, V12-, and V30 ROs**. D.** High-magnification images corroborate the presence of outer segments across all groups in the D210 dataset.

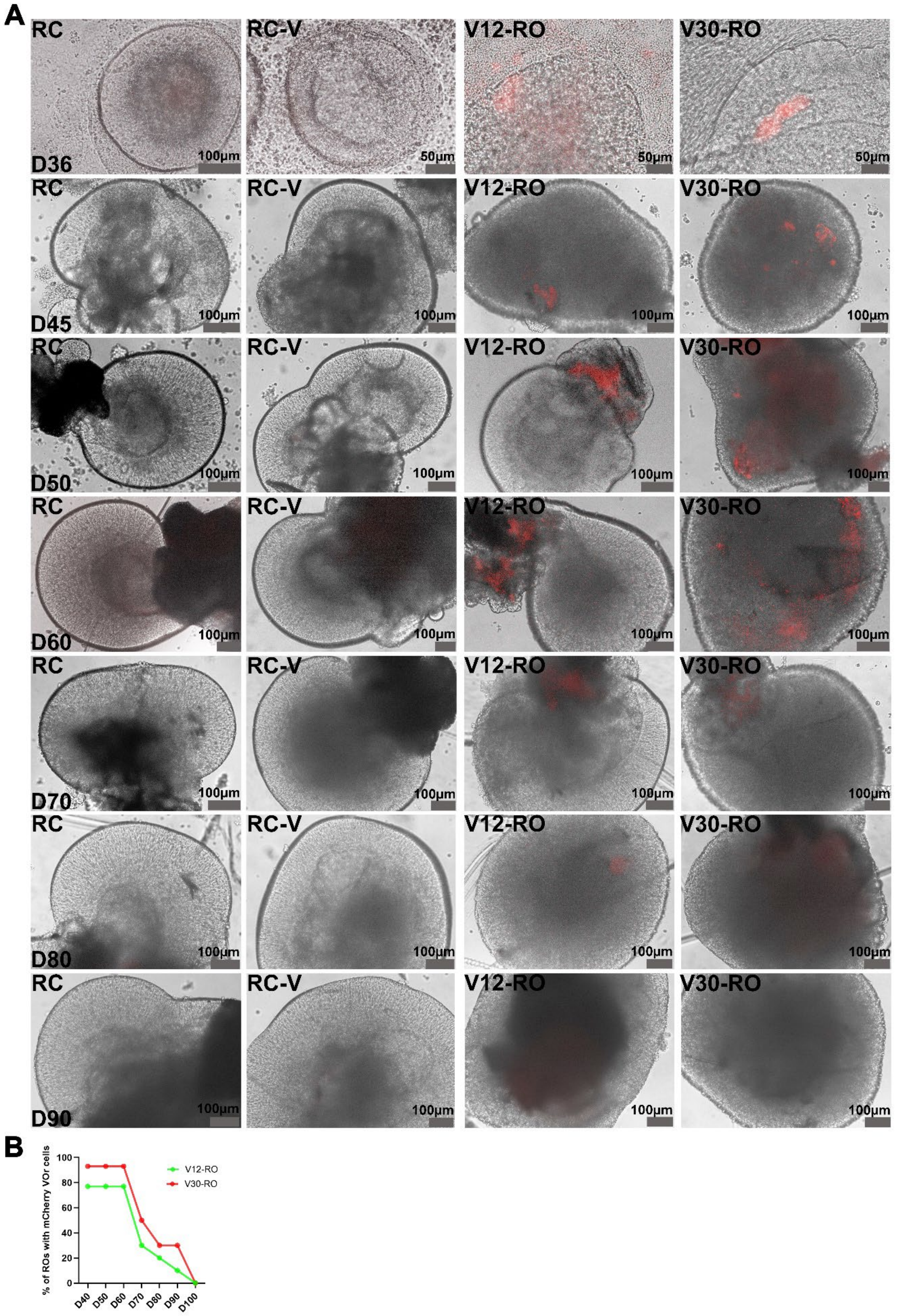


**Fig. S3: Transient presence of VOr-derived mCherry+ cells in developing ROs. A.** Representative merged bright-field and mCherry-fluorescence images showing VOr-derived mCherry$^+$ cells (red) in V12-RO and V30-RO cultures at the indicated RO differentiation days. mCherry$^+$ cells were observed near the developing retinal neuroepithelium following VOr-cell addition and became less readily detectable after D60. **B.** Quantification corroborated a decline in the percentage of $mCherry^+$ cells in V12 and V30 ROs between D60 and D100. Data are presented as mean percentage of $mCherry^+$ cells ± SEM (n=10).

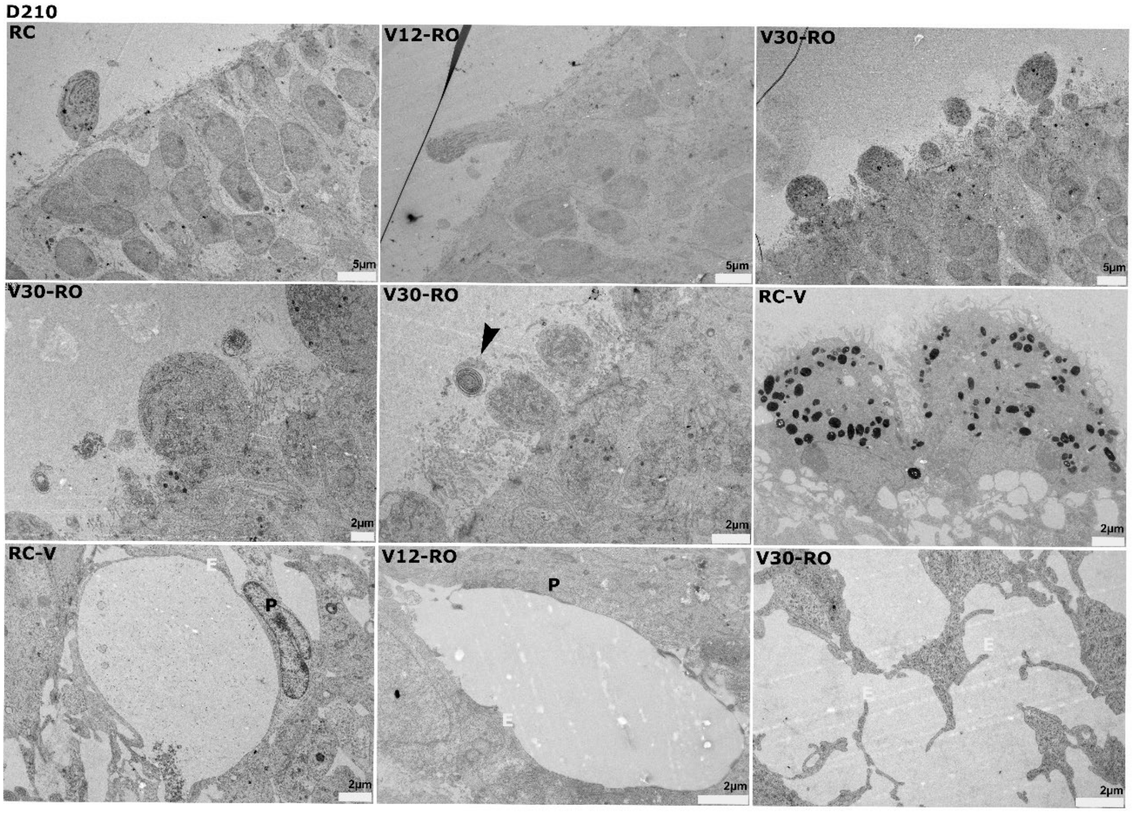


**Fig. S4: Transmission electron microscopy reveals vascular-like lumen structures in RC-V, V12- and V30 ROs at D210.** Transmission electron microscopy revealed lumen-like structures bordered by endothelial (E, white) and pericyte-like (P, black) cells in RC-V medium-control, V12- and V30 D210 ROs, consistent with the formation of vascular-like structures. RC-V ROs showed a high abundance of RPE. Black arrows mark photoreceptor outer segment-like structures.

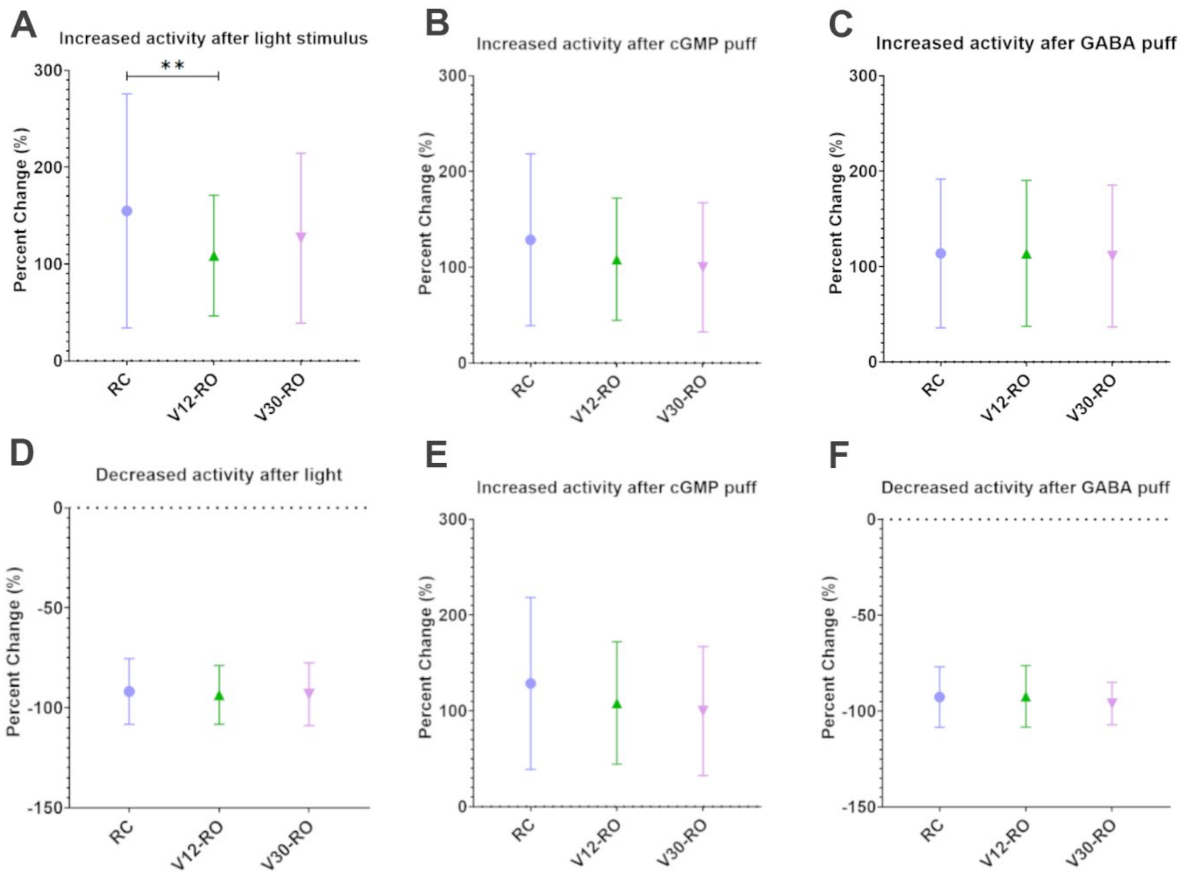


**Fig. S5: Functional assessment of ROs stimulated with light (A, D), cGMP (B, E), and GABA (C, F).** Graphs show the mean (+/- SD) percentage change of activity (spike counts) for RC-ROs (blue circle), V12-ROs (green triangle), and V30-ROs (purple inverted triangle) organoids. Number of RGCs in A) RC-ROs (165), V12-ROs (68) and V30-ROs (55), in B) RC-ROs (103), V12-ROs (74) and V30-ROs (34), in C) RC-ROs (166), V12-ROs (78) and V30-ROs (47), in D) RC-ROs (189), V12-ROs (144) and V30-ROs (109), in E) RC-ROs (248), V12-ROs (150) and V30-ROs (144) and in F) RC-ROs (209), V12-ROs (142) and V30-ROs (130). An unpaired T-test was used for the statistics: the p-value in A is 0.0031.

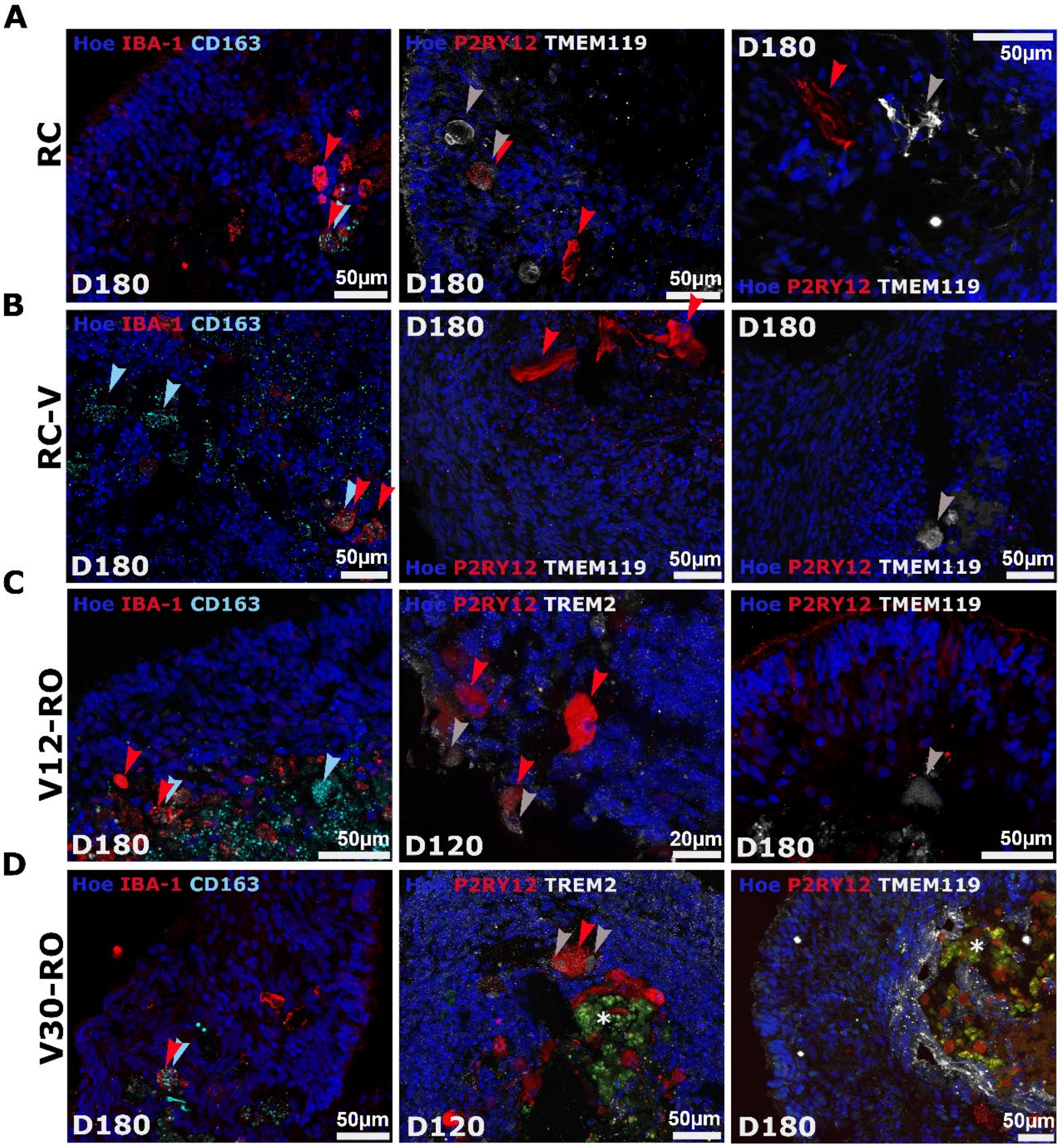


**Fig. S6: Immunofluorescence characterisation showed dynamic expression of putative microglia-like and BAM-like cells in all RO groups between D120 and D180.** IF characterisation of microglial expression reveals the presence of IBA1+ cells (red arrows) with both amoeboid and ramified morphologies at D180. Partial co-localisation of IBA1 with the CD163+ (light blue arrows) suggests the presence of BAMS across RO groups by D180. At day 120 of differentiation, V12- and V30-ROs show P2RY12+ (red)/TREM2+ (white) cells, consistent with a dynamic microglia-like phenotype and a subsequent loss of P2RY12+ in these groups by D180. Homeostatic ramified P2RY12+ morphologies are only observed in RC- and RC-V ROs in addition to a more ameboid P2RY12+/TMEM119+ transitioning phenotype also seen in RC ROs.

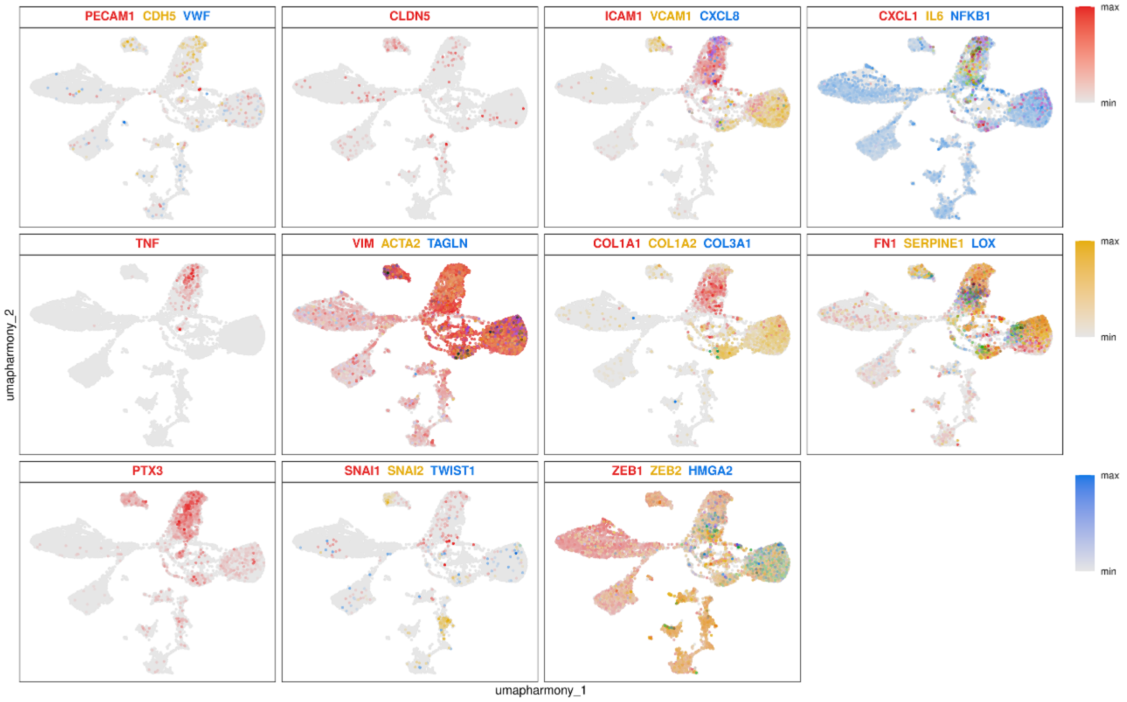


**Fig. S7: Expression of endothelial, inflammatory, and mesenchymal activation markers across the integrated single-cell transcriptomic landscape.** UMAP feature plots showing the expression patterns of endothelial-associated markers (*PECAM1, CDH5, VWF, CLDN5*), inflammatory and endothelial activation markers (*ICAM1, VCAM1, CXCL8, CXCL1, IL6, NFKB1, TNF*), and mesenchymal/perivascular markers (*VIM, ACTA2, TAGLN, COL1A1, COL1A2, COL3A1, FN1, SERPINE1, LOX, PTX3*). Expression of epithelial-to-mesenchymal transition (EMT)-associated transcription factors (*SNAI1, SNAI2, TWIST1, ZEB1, ZEB2*) and *HMGA2* is also shown. Each point represents an individual cell, with coloured points indicating relative transcript expression and grey points representing cells with low or undetected expression. UMAP coordinates are shown on the *x*- and *y*-axes.

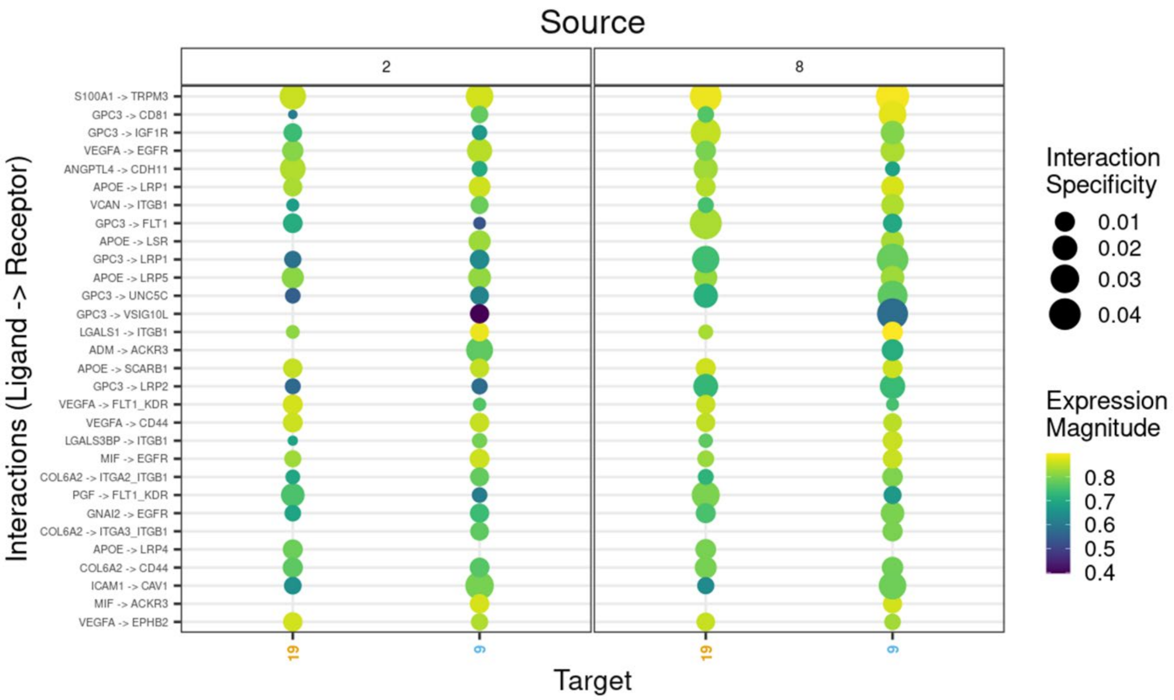


**Fig. S8: Predicted ligand–receptor interaction network between source and target cell populations.**

Bubble plot showing the top inferred intercellular communication events ranked by interaction specificity and expression magnitude. Each bubble represents a ligand–receptor pair, with bubble size and colour denoting the strength of the predicted interaction. Listed pairs include VEGFA–FLT1/KDR, VEGFA–EGFR, VEGFA–CD44, PGF–FLT1/KDR, APOE–LRP family interactions, GPC3-associated interactions, and extracellular matrix/adhesion-related signals such as VCAN–ITGB1, COL6A2–ITGA2/ITGB1, COL6A2–ITGA3/ITGB1, and ICAM1–CAV1. The source and target cell groups are indicated on the axes, with interaction specificity shown along the x-axis and expression magnitude along the y-axis.

A

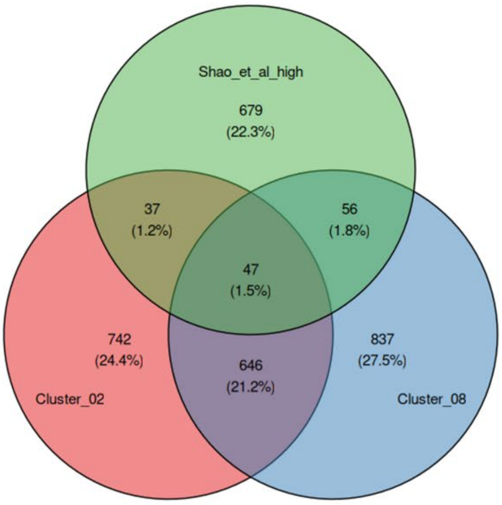


B

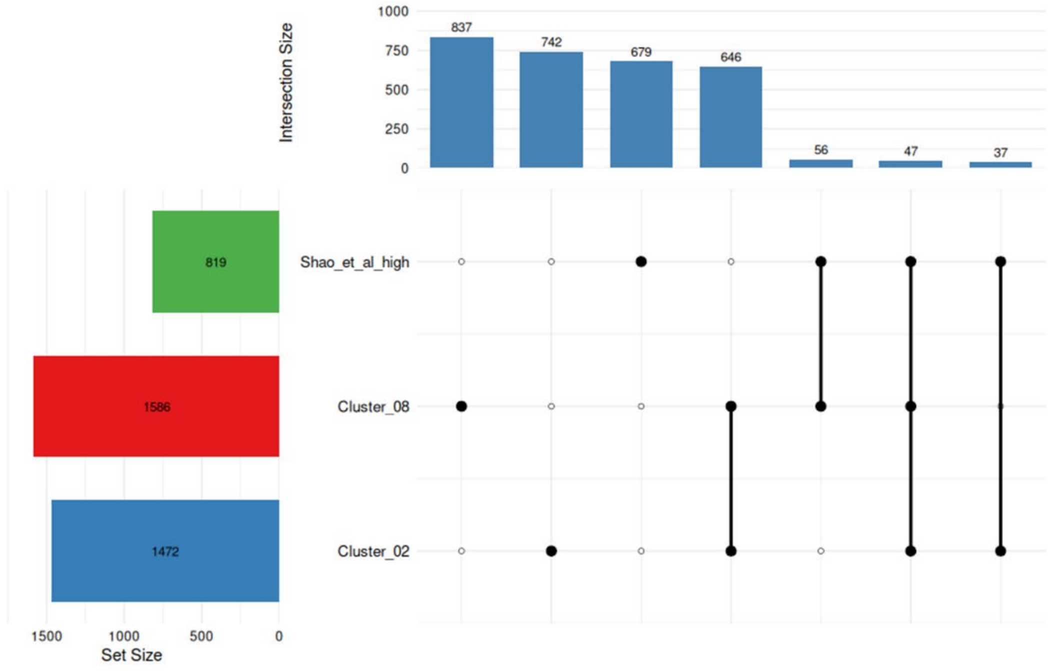


**Fig. S9: Transcriptomic comparison and gene set overlap between scRNA-seq clusters 2 and 8 and high-glucose-treated human retinal endothelial cells (hRECs). A.** Venn diagram displaying the overlap of differentially expressed gene sets among Cluster 2 (red), Cluster 8 (blue), and high-glucose-upregulated genes in hRECs from Shao et al. (GSE117238, green).

Numbers and percentages indicate unique and shared gene counts across the intersections. **B.** UpSet plot illustrating the specific set intersections and corresponding gene counts between Cluster 2, Cluster 8, and the high-glucose-associated gene signature. Representation-factor analysis demonstrated significant enrichment of high-glucose-induced genes specifically in Cluster 8 (RF $= 1.51$, $P = 1.60 \times 10^{-5}$), compared to non-significant enrichment in Cluster 2 (RF $= 1.32$, $P = 0.00485$), which did not reach the significance threshold of P < 0.001.
